# A diet-dependent enzyme from the human gut microbiome promotes Th17 cell accumulation and colitis

**DOI:** 10.1101/766899

**Authors:** Margaret Alexander, Qi Yan Ang, Renuka R. Nayak, Annamarie E. Bustion, Vaibhav Upadhyay, Katherine S. Pollard, Peter J. Turnbaugh

## Abstract

Aberrant activation of Th17 cells by the gut microbiota contributes to autoimmunity; however, the mechanisms responsible and their diet-dependence remain unclear. Here, we show that the autoimmune disease-associated gut Actinobacterium *Eggerthella lenta* increases intestinal Th17 cells and worsens colitis in a *Rorc*-dependent and strain-variable manner. A single genomic locus predicted Th17 accumulation. A gene within this locus, encoding the Cgr2 enzyme, was sufficient to increase Th17 cells. Levels of *cgr2* were increased in stool from patients with rheumatoid arthritis compared to healthy controls. Dietary arginine blocked *E. lenta*-induced Th17 cells and colitis. These results expand the mechanisms through which bacteria shape mucosal immunity and demonstrate the feasibility of dissecting the complex interactions between diet, the gut microbiota, and autoimmune disease.

**One Sentence Summary:** An autoimmune disease-associated bacterium triggers disease due to a diet-dependent enzyme that regulates mucosal immunity.

## Main Text

Intestinal immune responses are linked to the trillions of microorganisms that colonize the gastrointestinal tract (*1, 2*). Therefore, inter-individual variations in the gut microbiome could contribute to diseases associated with dysregulated immune responses such as autoimmunity (*3–5*). Consistent with this hypothesis, multiple disease models are ameliorated in germ-free (GF) or antibiotic-treated mice (*6–8*). Microbiome-wide association studies have identified bacteria enriched in autoimmune diseases, including rheumatoid arthritis (RA) (*3*), inflammatory bowel disease (IBD) (*4*), and multiple sclerosis (*5, 9*). However, whether or not these human gut bacterial strains play a causal role in disease remains largely unknown. Furthermore, the observation that host diet significantly impacts the structure and function of the gut microbiome (*10–12*) raises the question of whether or not diet-mediated shifts in the microbiome impact bacterial immune modulation and downstream disease phenotypes.

The mechanisms whereby the gut microbiome influences autoimmune disease and the role of diet in these interactions are even more mysterious. Perhaps the most well-studied mechanism is the bacterial-induced accumulation of T helper 17 (Th17) cells which are involved in maintaining the intestinal barrier and coordinating immune responses to extracellular pathogens and whose aberrant activation drives autoimmunity (*13*). Seminal work revealed that segmented filamentous bacteria (SFB) increase CD4+ IL-17a+ Th17 cells in mice (*14, 15*). Likewise, the composition and structure of the human gut microbial community significantly impact Th17 levels and severity of disease models following transfer into germ-free mice (*15, 16*). The prevalent human gut Actinobacterium *Bifidobacterium adolescentis*, which is enriched in IBD patients (*17*), also increases CD4+ IL-17a+ Th17 cells and worsens arthritis in a mouse model (*18*). Similar to SFB, the physical adhesion of *B. adolescentis* to the intestinal epithelium has been implicated in the mechanism of Th17 accumulation; however, it remains unclear if other gut bacterial species regulate Th17 cells through adhesion-independent mechanisms.

*Eggerthella lenta*, another prevalent gut Actinobacterium found in >80% of individuals (*19*), is enriched and coated with IgA in patients with autoimmune diseases (*3, 5, 20*), but the impact of this bacterium on host immunity and disease phenotypes remains unexplored. Here, we show that *E. lenta* colonization is sufficient to increase intestinal CD4+ IL-17a+ (Th17) cell levels and exacerbates murine colitis. Through a combination of gnotobiotics, comparative genomics, *in vitro* assays, and bacterial genetics we identify a single strain-variable gene (cardiac glycoside reductase 2, *cgr2*) which is sufficient to increase Th17 cells. These data provide the first physiological function for Cgr2, which was previously implicated in xenobiotic metabolism, inactivating the cardiac drug digoxin and other toxic plant compounds within the cardenolide family (*12, 21*). *Cgr2* copy number is significantly higher in RA patient stool samples than healthy controls, supporting the translational relevance of these findings. Finally, by leveraging the knowledge that Cgr2 activity is abrogated by high levels of dietary arginine (*12, 21*) we show that dietary arginine prevents *E. lenta*-dependent Th17 accumulation and associated colitis emphasizing that diet can affect the capacity of specific bacterial strains to modulate immune responses and downstream disease.

*E. lenta* colonization of germ-free (GF) mice increases Th17 activation. GF mice colonized with *E. lenta* strain UCSF 2243 for two weeks had increased percentages of IL-17a+ CD4+ (Th17) cells within the CD3+ population in the small intestine and colon but not spleen (Figs. 1A-F, fig. S1A-B). Increases in IL-17a+ CD4+ percentages were accompanied by increased IL-17a mean fluorescence intensity (MFI) in the small intestine and colon (Figs. 1C and F). However, levels of CD4+ Rorγt+ cells and Rorγt MFI were not altered in *E. lenta* colonized mice in any of these tissues compared to GF mice (Figs. 1G-L, figs. S1C-D). *E. lenta* monocolonization did not have a significant effect on intestinal IFNγ+ CD4+ T cells, IL17a+ TCRγδ+ T cells, or lineage-Rorγt+ IL-17a+ group 3 innate lymphoid cells (ILC3) compared to GF mice (figs. S1E-L). However, while lineage-Rorγt+ IL-22+ cells were decreased in *E. lenta* monocolonized mice, total IL-22 MFI was not altered in *E. lenta* monocolonized mice, suggesting that IL-22 producing ILC3 cells are decreased in *E. lenta* monocolonized mice but total expression of IL-22 is not changed (figs. S1K, and S1M-N). qPCR of GF and *E. lenta* monocolonizaerd mice ileal tissue revealed an increase in Rorγt targets IL-17f, CCR6, a trending increase in IL-23r, but no significant differences in IL-22 and Rorγt levels (Fig. 1M). There were no significant changes in Th1 associated transcripts (IFNγ, Tbet), Treg associated transcript (FoxP3), or other inflammatory markers, with the exception of a trend towards increased BCL6, which could indicate an increase in T follicular helper cells (Fig. 1M and fig. S1O). To test if *E. lenta* promotes IL-17a+ CD4+ accumulation in the context of a more complex microbiota, we gavaged SPF mice with *E. lenta* or a media control (BHI) every other day for two weeks and observed a similar accumulation of IL-17a+ CD4+ cells in the small intestinal and colonic lamina propria and increased IL-17a MFI with *E. lenta* (Fig. 1N-S). We observed a similar pattern in the CD4+ IL-17f+ population and IL-17f MFI but again no significant differences in the CD4+ Rorγt+ population (figs. S2A-H). We further tested whether secreted factors from *E. lenta* were sufficient to promote Th17 accumulation by gavaging SPF mice with conditioned media from *E. lenta* or a BHI control and observed a similar increase in CD4+ IL-17a+ cells, IL-17a MFI, and IL-17f MFI in the small intestine (figs. S2I-L).

**Fig. 1.**
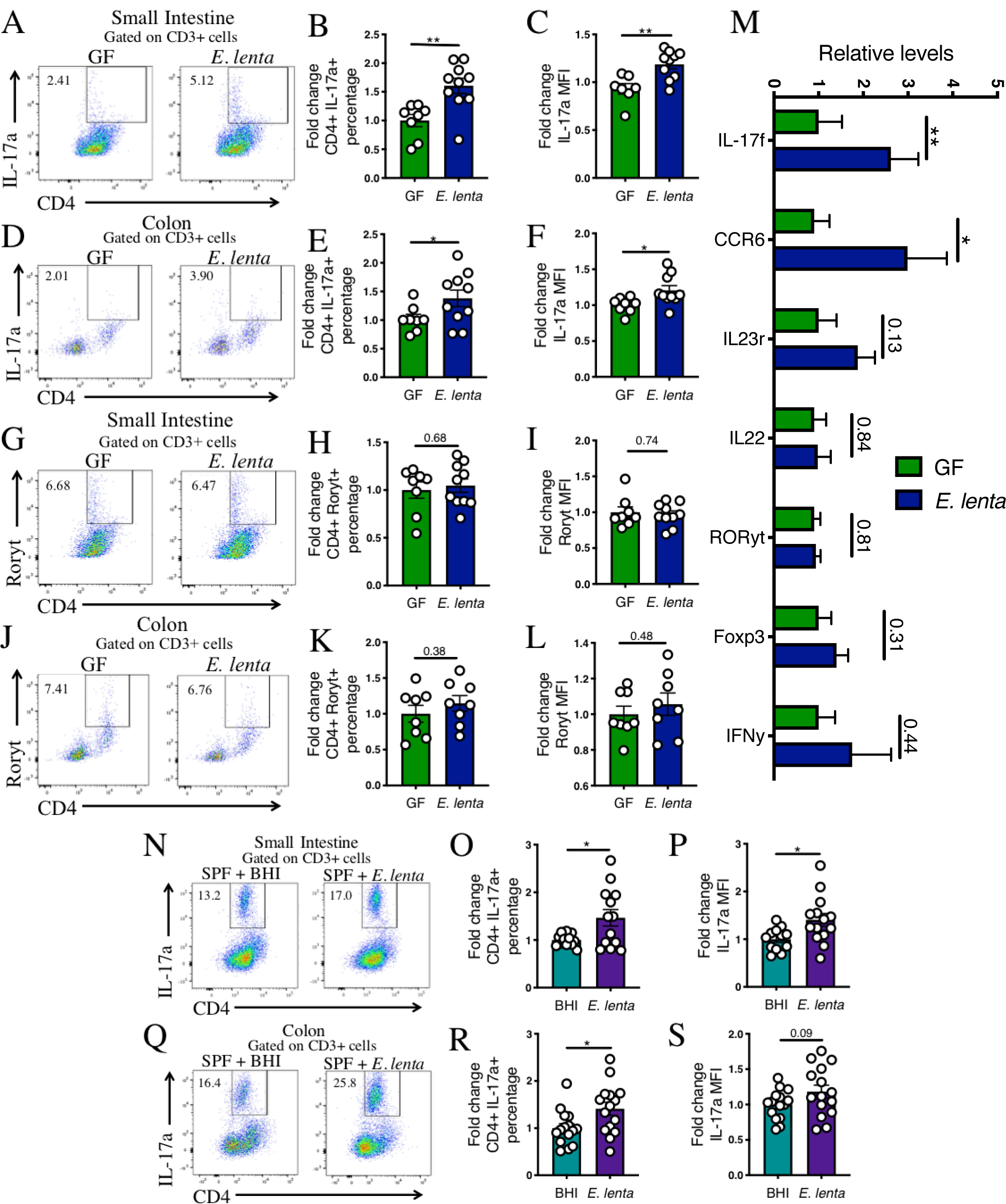
*E. lenta* colonization increases intestinal CD4+ IL-17a+ cells in the presence and absence of an intact gut microbiota. (A-F) IL-17a+ CD4+ Th17 levels within the CD3+ population and IL-17a mean fluorescence intensity (MFI) were measured via flow cytometry in the small intestinal and colonic lamina propria of germ-free (GF) mice or mice monocolonized with *E. lenta* strain UCSF 2243 for two weeks (n=8-10). Representative flow plots of the IL-17a+ CD4+ population within the CD3+ gate are displayed and fold changes (relative to GF) of this population and IL-17a MFI are quantified to the right. Data are pooled from two independent experiments with each point representing an individual mouse. (G-L) Representative flow and fold change (relative to GF) of CD4+ Rorγt+ cells within the CD3+ population and MFI of Rorγt in the small intestinal and colonic lamina propria from the same mice in A-F. (M) RT-qPCR panel of immune genes from GF or *E. lenta* monocolonized ileal samples (n=10-16). Data are relative to GF. (N-S) C57BL/6J SPF mice were gavaged every other day for 2 weeks with BHI media control or *E. lenta* strain 2243 (n=10-11). Representative flow plots of (N) small intestinal and (Q) colonic lamina propria IL-17a+ CD4+ cells within the CD3+ population and fold change of this population and IL-17a MFI are plotted to the right. Data are from three independent experiments. Each point represents an individual mouse and the mean±SEM is plotted. \**P* < 0.05; \*\**P* < 0.01; or listed, Welch’s t-test. See figs. S1, S2, and S3 for gating strategy and additional related data.

Prior studies have found *E. lenta* at higher levels in patients with rheumatoid arthritis (*3*) and multiple sclerosis (*5, 9*) relative to healthy controls, suggesting a general association between this bacterial species and autoimmune disease. To assess the broader relevance of *E. lenta* in human autoimmune disease, we analyzed two metagenomic datasets totaling 105 IBD patients (52 ulcerative colitis and 53 Crohn’s disease) and 100 healthy subjects (*22–24*). A combined analysis revealed that *E. lenta* was significantly higher in IBD patient stool samples relative to healthy controls (figs. S4A-F). These results indicate that there may be a general mechanism linking *E. lenta* to multiple autoimmune diseases.

Next, we turned to mouse models of IBD due to the ability to readily use these models in a gnotobiotic system and associations with *E. lenta* levels, allowing us to test the impact of *E. lenta* colonization on the severity of autoimmune disease phenotypes. GF mice were colonized 2 weeks with *E. lenta* prior to DSS colitis induction to assess the impact of Th17 activation on disease phenotypes. *E. lenta* colonized mice developed more severe disease than their GF counterparts in both disease scores and colon shortening, crypt destruction as observed via histology, elevated lipocalin levels, and increased colonic Th17 levels (Figs. 2A-D and figs. S5A-D). To determine if *E. lenta* mediated Th17 accumulation contributed to this worsened disease phenotype, we compared *WT* and *Rorc*^*-/-*^ SPF mice gavaged with *E. lenta* or BHI media control every other day for 2 weeks preceding induction of DSS colitis. *E. lenta* colonized *WT* SPF mice developed more severe disease compared to BHI controls while *Rorc*^*-/-*^ mice had no differences in disease severity between the BHI and *E. lenta* groups (Figs. 2E-H). Th17 levels were elevated in the colon of *WT* mice gavaged with *E. lenta* but, as expected, there were low levels of these cells and no difference between *E. lenta* gavage and control in the *Rorc-/-* mice (figs. S5E-H). These results suggest that Th17 accumulation by *E. lenta* contributes to worsened colitis severity in *E. lenta* colonized mice, although it is important to note that Th17 cells are not the only cell type missing from *Rorc*^*-/-*^ mice as ILC3 cells are also lacking and these mice additionally have issues with thymopoiesis (*25*). To test whether *E. lenta* colonization contributes to more severe disease in an independent model of colitis, we gavaged *E. lenta* or a media control (BHI) three times a week into *IL-10*^*-/-*^ SPF mice and tracked weights and survival. *E. lenta* gavage resulted in lower percentage of initial weight, decreased survival, and increased lipocalin compared to the media control (Figs. 2I-K) suggesting that *E. lenta* colonization contributes to more severe disease in a chronic colitis model.

**Fig. 2.**
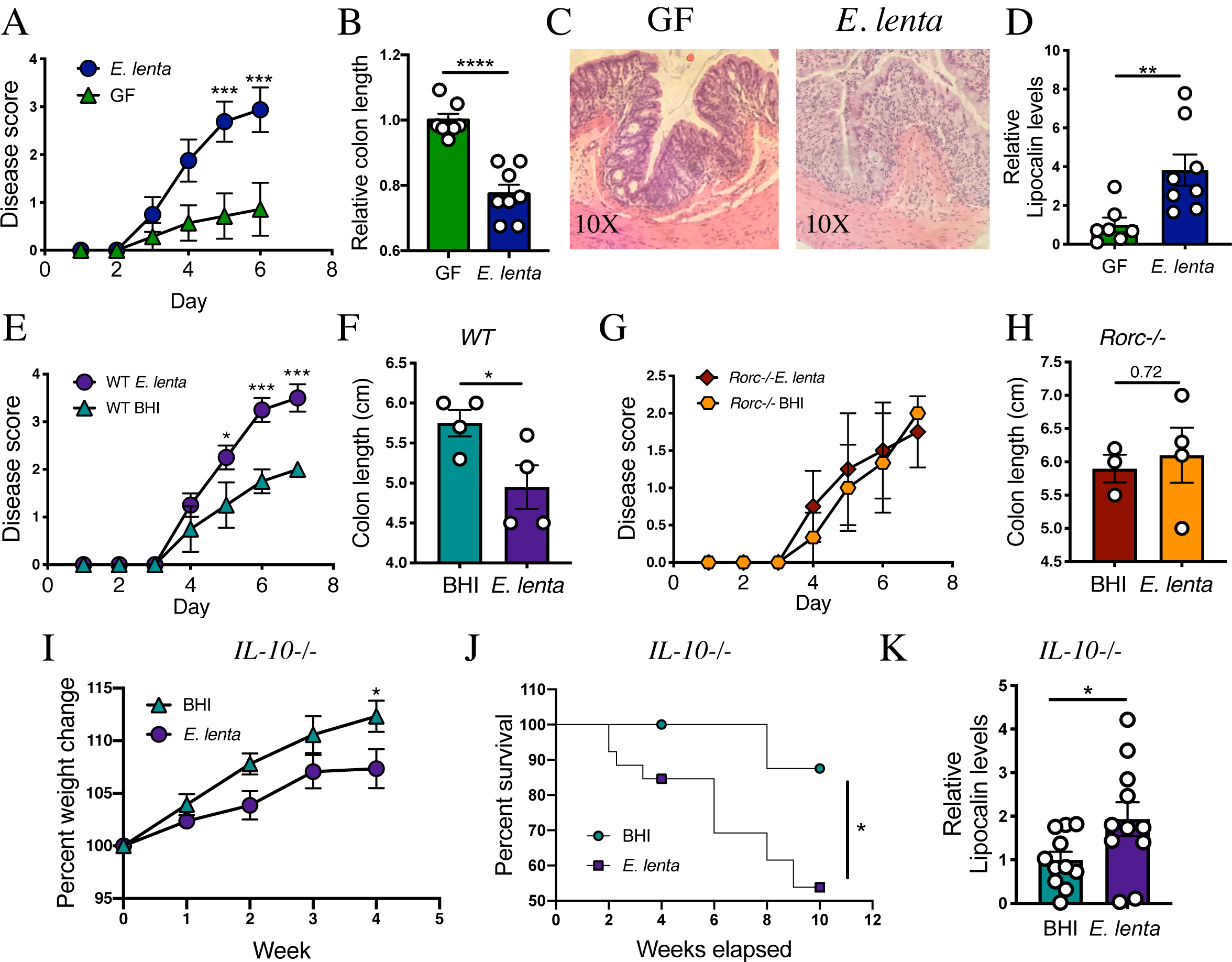
*E. lenta* colonization contributes to worsened colitis in a Rorc-dependent manner. (A-D) Germ-free (GF) or *E. lenta* strain 2243 monocolonized mice were treated with 2% DSS for 6 days and (A) disease was scored based on published scoring metrics (*41*). \*\*\**P* < 0.001 two-way ANOVA with Sidak’s multiple comparison correction. Data are from 2 independent experiments (n=7-8). (B) Colon shortening was assayed and relative colon length is displayed (relative to GF). \*\*\*\**P* < 0.0001 (Welch’s t-test). (C) Representative histology from GF or *E. lenta* monocolonized colons with H&E staining at 10X magnification. (D) Relative lipocalin levels as measured via ELISA from the colon content. \*\**P* < 0.01 (Welch’s t-test) (E-H) SPF *WT* or *Rorc*^*-/-*^ mice were gavaged with a BHI media control or *E. lenta* 2243 every other day for 2 weeks then treated with 2% DSS for 7 days and disease was tracked. (E) Disease scores over time in *WT* SPF mice treated with BHI or *E. lenta.* \**P* < 0.05, \*\*\**P* < 0.001 two-way ANOVA with Sidak’s multiple comparison correction (n=4). (F) Day 7 colon lengths in the same mice as E. (G) Disease scores over time in *Rorc-/-* SPF mice treated with BHI or *E. lenta.* (H) Day 7 colon lengths in the same mice as G. *P* value stated in Welch’s t-test. Data are from 1 representative experiment. (I-K) *IL-10*^*-/-*^ mice were gavaged with a BHI media control or *E. lenta* 2243 three times a week for 6 weeks and (I) percentage of initial weight was tracked (n=11). \**P* < 0.05 two-way ANOVA with Sidak’s multiple comparison correction. Mice that dropped out of the study before week 4 were removed from weight change analysis. (J) Percent survival of *IL-10-/-* mice gavaged with BHI or *E. lenta* 2243 three times a week for 10 weeks (n=19 (BHI) 26 (*E. lenta*)). Mice were sacrificed when rectal prolapse developed. Data are from 2 independent experiments. \**P* < 0.05 Log-rank Mantel-Cox test. (K) Lipocalin levels were measured via ELISA from the colon content after 6-10 weeks of gavage. \**P* < 0.05 (Welch’s t-test). Data are from 2 independent experiments and set relative to BHI (n=11). Mean±SEM is displayed. Each dot represents an individual mouse. See also fig. S5 and data S1 for related data.

*E. lenta* mediated Th17 accumulation is a strain-variable phenotype. We monocolonized mice with *E. lenta* strains UCSF 2243 (2243), FAA 1-3-56 (1356), DSM 15644 (15644), AB12#2 (AB12), or heat-killed *E. lenta* 2243 (see table S1 for strain information). Viable *E. lenta* 2243 and AB12 were the only groups which elevated Th17 levels compared to GF mice in intestinal compartments (Figs. 3A-D and figs. S6A-C) indicating *E. lenta* mediated Th17 accumulation is strain variable and requires live bacteria. Interestingly, the AB12 strain promoted Th17 accumulation to a greater extent in the colon than the small intestine compared to the 2243 strain which promoted Th17 accumulation in both the small intestine and colon. Next, we tested whether monocolonization with a strain of *E. lenta* that did not increase Th17 levels would impact DSS colitis. While monocolonization with *E. lenta* 2243 caused more severe colitis, colonization with *E. lenta* 1356 did not, as observed with lower disease scores, less severe colon shortening, lower colon lipocalin levels, and fewer intestinal Th17 cells (Figs. 3E-G, fig. S6D-F).

**Fig. 3.**
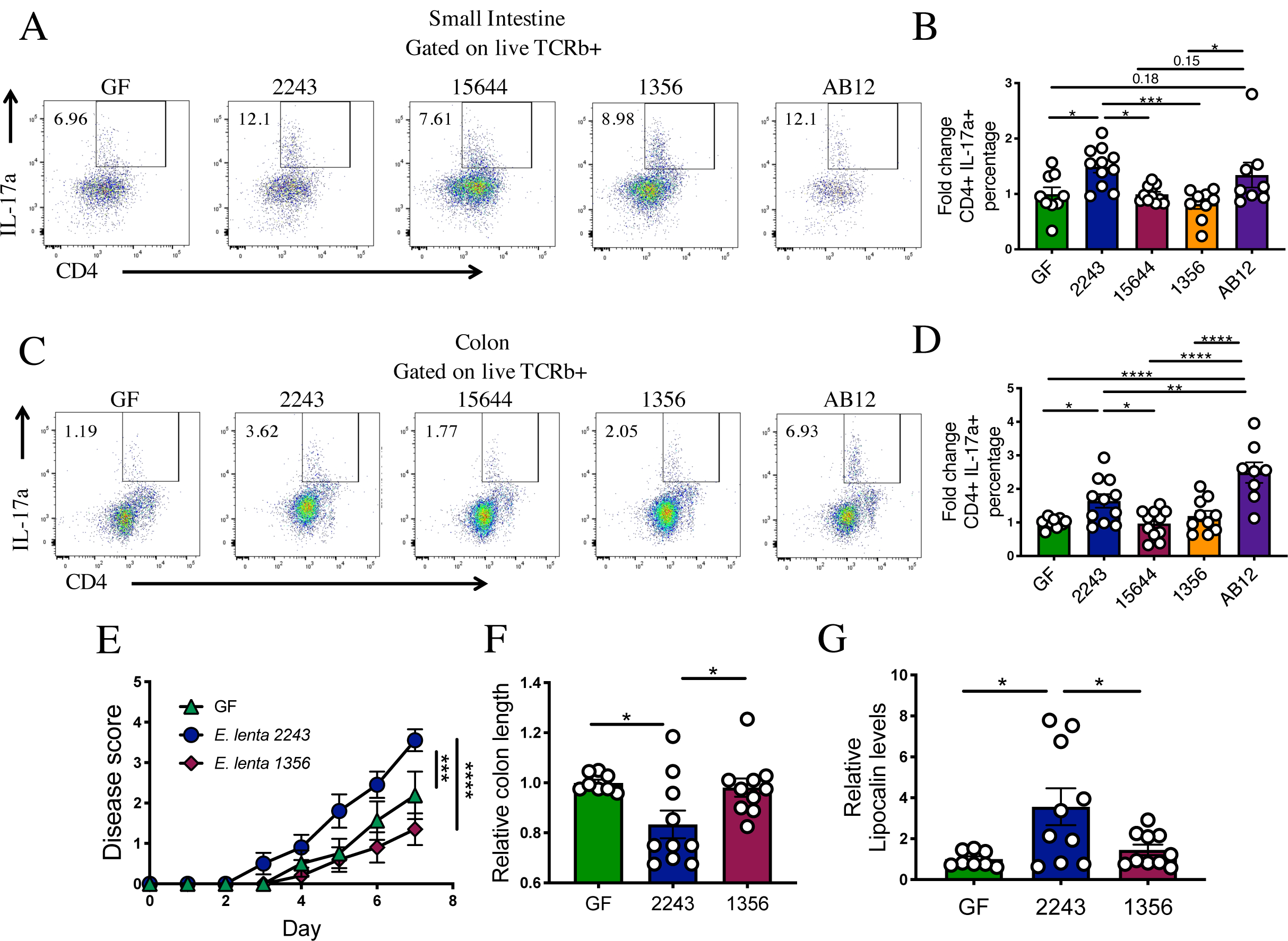
Th17 accumulation and colitis severity varies between *E. lenta* strains. (A-D) Th17 levels were analyzed in germ-free (GF) mice or mice monocolonized with *E. lenta* 2243 (2243), *E. lenta* 1356 (1356), *E. lenta* 15644 (15644), or *E. lenta* AB12#2 (AB12) as measured by IL-17a+ CD4+ within the live TCRb+ population in the (A-B) small intestinal and (C-D) colonic lamina propria. Representative flow plots are shown and fold change compared to the GF group is quantified to the right. Data are from 2 independent experiments (n=8-11). (E-I) GF, mice or monocolonized with *E. lenta* 2243, or *E. lenta* 1356 for 2 weeks were treated with 2% DSS and (E) disease was tracked for 7 days (n=8-10). \*\*\**P* < 0.001; \*\*\*\**P* < 0.0001 two-way ANOVA with Sidak’s multiple comparison correction. (F) Day 7 colon lengths. (G) Lipocalin levels as quantified via ELISA in the colon content. \**P* < 0.05; \*\**P* < 0.01; \*\*\**P* < 0.001 (one-way ANOVA with Tukey multiple comparison test unless otherwise noted). Mean±SEM is displayed. Each dot represents an individual mouse. See figs. S3 for gating strategies and S6, table S1, and Data S1 for related data.

To assess the variation between *E. lenta* strains and their capacity to increase Th17 cells, we turned to an *in vitro* cell culture assay. Splenic CD4+ T cells were cultured in Th17 skewing conditions with the addition of conditioned media from *E. lenta* strains or media controls. Recapitulating our *in vivo* findings, conditioned media from *E. lenta* 2243 but not 1356 or 15644 significantly increased IL-17a levels compared to media control as assessed by IL-17a ELISA (Fig. 4A). Of note, unconditioned media (BHI) decreased IL-17a production, suggesting there is a factor in our bacterial culture media inhibiting its production. With our *in vitro* system, we wanted to further characterize the mechanism by which *E. lenta* strain 2243 conditioned media alters IL-17a levels. Heat treatment or filtering through a 3 kDa filter did not abrogate the IL-17a inducing activity of *E. lenta* strain 2243 conditioned media compared to BHI controls (fig. S7A) suggesting a small molecule is involved in IL-17a induction in our *in vitro* setting. Further, we assessed whether the conditioned media was changing Th17 differentiation, proliferation, or expression of IL-17a by altering the timing of conditioned media addition (fig. S7B). *E. lenta* strain 2243 conditioned media resulted in higher IL-17a levels compared to controls whether added at the time of Th17 skewing or after skewing had already occurred (figs. S7C and 7D). However, proliferation levels were not different between conditions (figs. S7E and 7F). These results suggest that the 2243 conditioned media can impact IL-17a production after cells have been differentiated, which supports a mechanism where *E. lenta* 2243 acts upon IL-17a production, not Th17 proliferation.

**Fig. 4.**
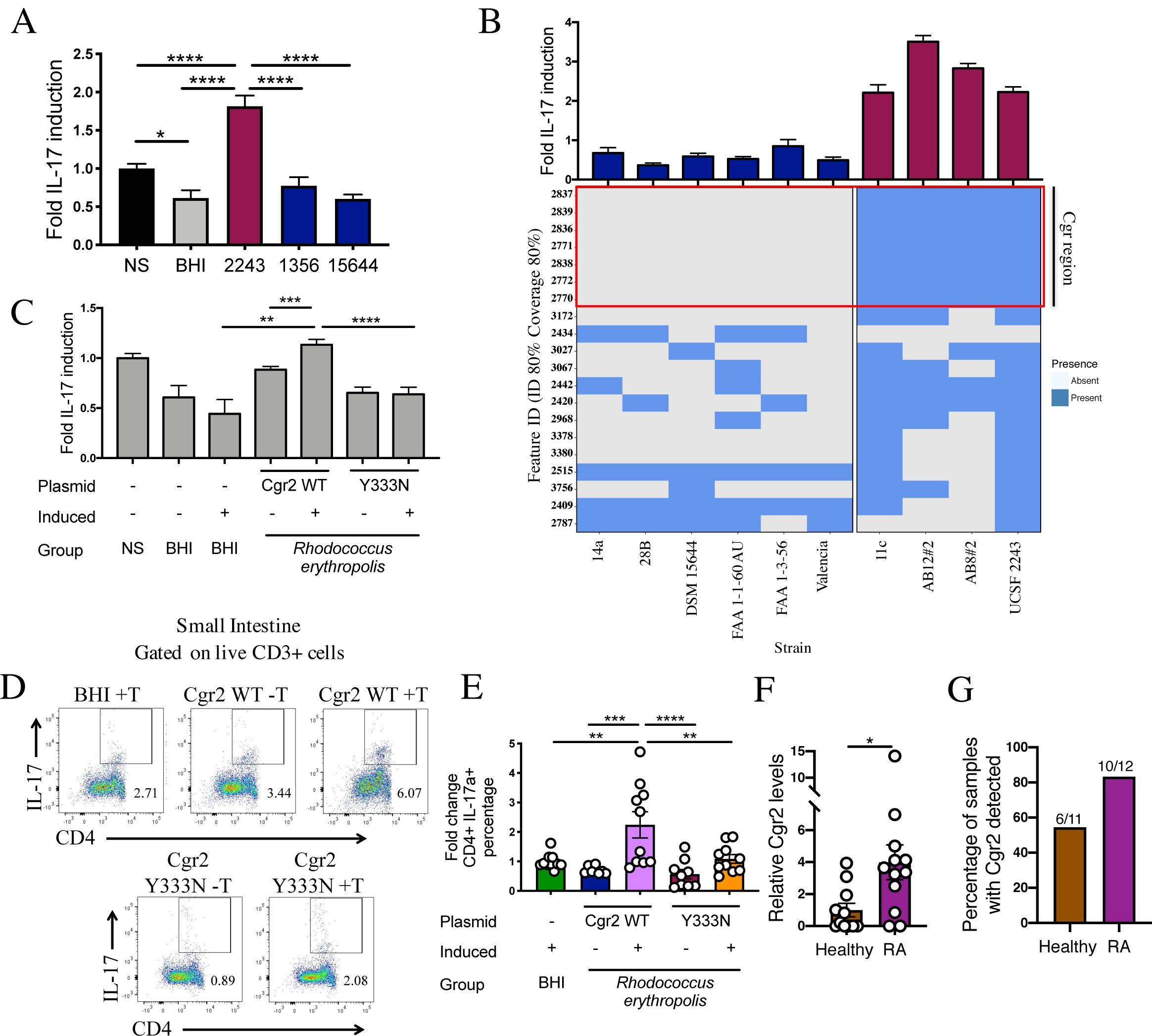
A single strain-variable genomic locus is predictive and sufficient for IL-17a induction. (A) Splenic T cells were treated with Th17 skewing conditions and conditioned media from *E. lenta* strains (2243, 1356, or 15644), a BHI media control, or no treatment (NS). IL-17a levels in the supernatant were measured via ELISA after 4 days of skewing followed by overnight PMA ionomycin restimulation (n=18-20 biological replicates (wells)/group). Levels of IL-17a are relative to the no treatment control which is set to 1. (B) In the same experimental design as A, 10 strains of *E. lenta* were screened and levels of IL-17a are relative to the no treatment control which is set to 1. Strains were classified into non-IL-17a inducers (fold IL-17a levels below 1) or high (fold IL-17a levels above 1). With these classifications we performed comparative genomics using ElenMatchR with 80% coverage and 80% minimum identity (*19*) to determine genomic regions shared between the 4 inducing strains and absent from the 6 non-inducing strains (n=8-12 biological replicates (wells)/group). Feature ID numbers 2770-2839 correspond to the cardiac glycoside reductase (*cgr*)-associated gene cluster (*21*) which is highlighted by the red box. (C) With our *in vitro* Th17 system we tested *Rhodococcus erythropolis* stains with induced (+Thiostrepton) or uninduced (-Thiostrepton) expression of WT Cgr2 (n=14), a natural partial loss-of-function variant Y333N Cgr2 (n=12) with no treatment (NS) (n = 14), BHI (+/- Thiostrepton) (n=12 and n=8) as controls. Levels of IL-17a are relative to the no treatment control. \*\**P* < 0.01; \*\*\**P* < 0.001; \*\*\*\**P* < 0.0001 Welch’s t-test. (D-E) C57BL/6J SPF mice were gavaged with conditioned media from *R. erythropolis* strains with induced (+Thiostrepton) or uninduced (-Thiostrepton) expression of WT Cgr2, Y333N Cgr2, or a BHI media control (with Thiostrepton). Representative flow plots of CD4+ IL-17a+ within the live CD3+ population in the small intestinal lamina propria lymphocytes are displayed and fold change relative to the BHI group quantified to the right (n=10). Each point represents an individual mouse. Data represents a combination of at least two independent experiments for A-E. (F) Relative levels of *cgr2* as measured via qRT-PCR from fecal samples from rheumatoid arthritis (RA) patients or healthy controls before therapeutic intervention and normalized for input mg of sample. Levels are relative to the healthy group (n=11-12). \**P* < 0.05 (Mann-Whitney test). (G) The percentage of RA or healthy stool samples with detectable *cgr2*. Numbers listed are number positive over total samples in the group. Mean±SEM is displayed. \**P* < 0.05; \*\**P* < 0.01; \*\*\**P* < 0.001; \*\*\*\**P* < 0.0001 (one-way ANOVA with Tukey multiple comparison test unless otherwise noted). Mean±SEM is displayed. Each dot represents an individual mouse or human. See table S1 for strain metadata. Raw values for IL-17a ELISA can be found in table S2. See figs. S3 for gating strategies and S7 and S8 for related data.

Having validated our *in vitro* assay, we tested a broader panel of 10 *E. lenta* strains (*19*) for their IL-17a induction capacity. This screen revealed 6 *E. lenta* strains that did not increase IL-17a production and 4 strains that significantly increased IL-17a production (Fig. 4B). Using the ElenMatchR comparative genomics tool (*19*), we determined that a single genomic locus (orthologous gene cluster 2770-2839), was present in all of the strains that increased IL-17a production and missing from all of the strains that did not induce IL-17a (Fig. 4B, figs. S8B-D). We had previously implicated this same locus, referred to as the cardiac glycoside reductase (*cgr*)- associated gene cluster, in the metabolism of the cardiac drug digoxin and other plant toxins from the cardenolide family (*21*). These results indicated that one or more of the genes in this locus also plays a role in shaping host immunity.

Based on prior evidence that digoxin inhibits Th17 cell differentiation (*26, 27*), we hypothesized that the enzyme sufficient for digoxin reduction, Cgr2, may be capable of increasing Th17 cells due to the reduction of an endogenous substrate found in the mouse gut lumen and our BHI cell culture media. A key prediction of this hypothesis is that Cgr2 should be sufficient to promote IL-17a production in our *in vitro* assay. We used constructs in which WT *cgr2* or a partial loss-of-function natural variant of *cgr2* (Y333N) were expressed in *Rhodococcus erythropolis* under control of a thiostrepton-inducible promoter (*21*). Expression of the WT but not variant form of *cgr2* resulted in a significant increase in IL-17a levels compared to uninduced controls (Fig. 4C). To confirm that *cgr2* expression is also sufficient to increase Th17 cells *in vivo*, we supplied SPF mice with conditioned media from *R. erythropolis* strains with WT or Y333N *cgr2* in the presence or absence of the thiostrepton inducer. Conditioned media from *R. erythropolis* with induced WT *cgr2* expression resulted in a significant increase in small intestinal Th17 cells compared (Fig. 4D and 4E). The partial loss of function mutant Y333N displayed an intermediate phenotype where Th17 levels were increased compared to uninduced controls but to a lesser magnitude than the WT *cgr2* version (figs. S8E-G).

Due to our findings that *cgr2* is sufficient to induce Th17 cells we postulated that the *cgr2* gene locus may be associated with Th17 driven diseases such as rheumatoid arthritis (RA). We analyzed the levels of *cgr2* via qPCR from the stool of patients with RA before any therapeutic intervention and healthy controls and found that *cgr2* levels were significantly increased in RA patient stool; 83% (10/12) of the RA patient samples had detectable *cgr2* compared to 54% (6/11) in healthy controls (Fig. 4F-G; *p*<0.05, one-sided Fisher’s exact test). These results provide initial support for the translational relevance of this specific bacterial gene for human autoimmune diseases.

Prior studies have shown that elevated protein intake, likely due to the increased concentration of arginine in the gut lumen, inhibits the ability of Cgr2 to reduce the cardiac drug digoxin (*12, 21*). We hypothesized that arginine would have a similar ability to prevent *E. lenta* mediated Th17 accumulation and associated disease phenotypes. GF, *E. lenta* strain 2243 (a *cgr+* strain), or 15644 (a *cgr-* strain) monocolonized mice were fed a 1% or 3% arginine diet. *E. lenta* 2243 colonized mice had elevated Th17 cells on the 1% arginine diet, but those levels were decreased back to baseline levels on the 3% arginine diet (Figs. 5A and 5B), and the different levels of Arg did not have a significant impact on the level of *E. lenta* colonization (fig. S9A). Consistent with our hypothesis, Th17 cells were significantly decreased on 3% arginine diet in *cgr+* 2243, whereas the opposite was seen in *cgr-* 15644 although to a lesser magnitude than the effect of strain 2243 (figs. S9C-D). There was no impact of arginine levels in GF mice (fig. S9B). As an additional control, we monocolonized mice on a 1% or 3% arginine diet with *Bifidobacterium adolescentis*, which has been shown to induce Th17 cells (*18*). *B. adolescentis* BD1 showed a similar response to arginine as *E. lenta* strain 15644, where the 3% arginine diet condition had trending higher levels of Th17 cells compared to the 1% arginine diet (fig. S9E). These results suggest that dietary arginine can have opposing effects on Th17 cells depending on gut bacterial colonization; higher arginine decreases Th17 levels in the context of *E. lenta* strain 2243 colonization but increases Th17 levels in mice colonized with *E. lenta* strain 15644 or *B. adolescentis.* To determine if the effect of arginine levels on *E. lenta* mediated Th17 accumulation alters colitis outcome, we gavaged *E. lenta* 2243 (*cgr+*), or 15644 (*cgr-*) into SPF mice on a 1% or 3% arginine diet every other day for two weeks and then induced DSS colitis. Mice gavaged with *E. lenta* 2243 (*cgr+*) on a high arginine diet had decreased Th17 cells and reduced disease severity compared to those on a low arginine diet (Figs. 5C-G). Differing arginine levels had no significant effects on disease severity or Th17 levels in the BHI or *E. lenta* strain 15644 (*cgr-*) treated groups (Figs. 5C-G). These results demonstrate that, in the context of colonization with a *cgr+ E. lenta* strain, increased arginine in the diet decreases both Th17 accumulation and colitis severity.

**Fig. 5.**
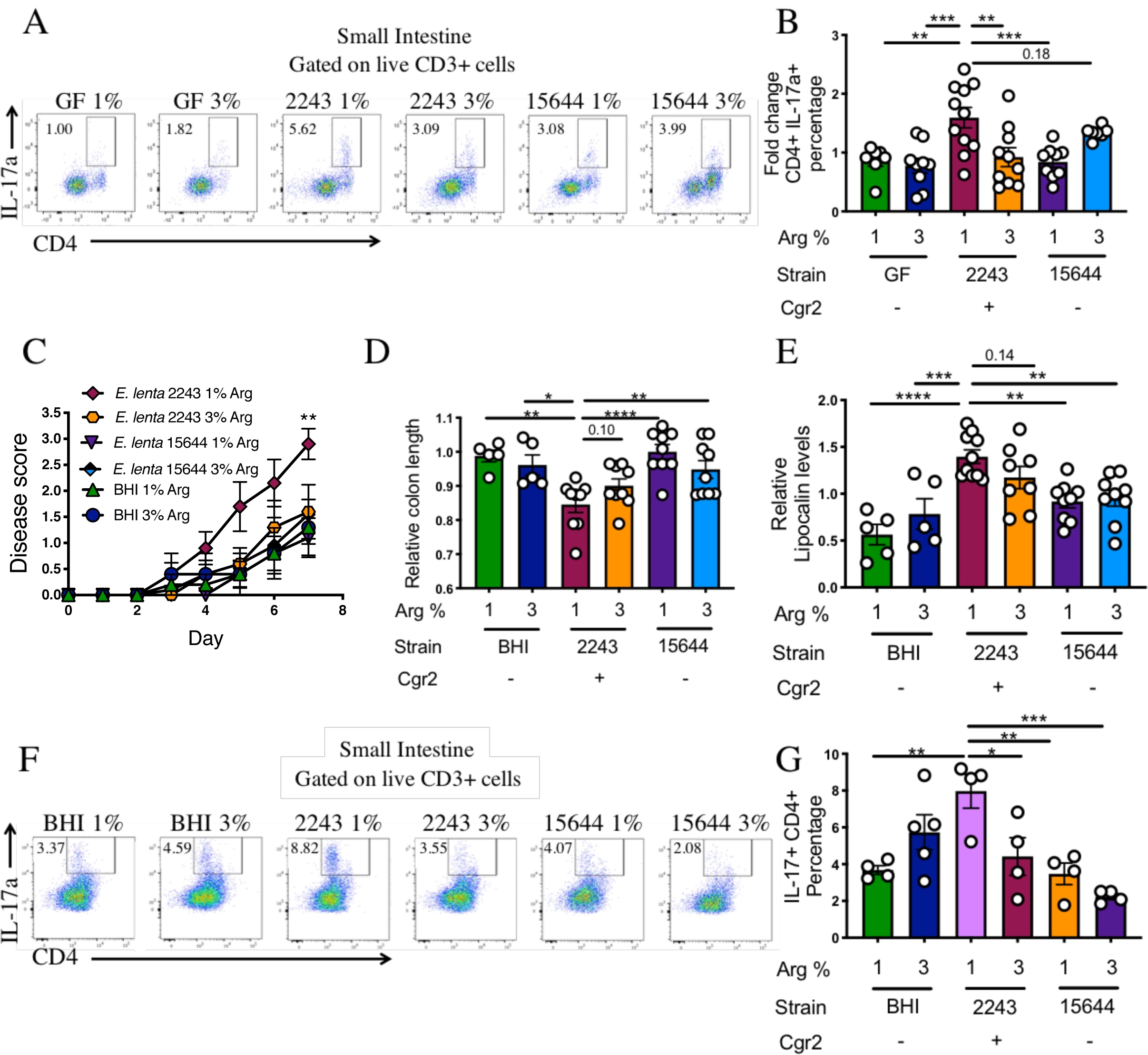
A high arginine diet inhibits Th17 accumulation and colitis induction by a *cgr+ E. lenta* strain. (A-B) Mice on a 1% or 3% arginine diet either remained germ-free (GF) or were monocolonized with *E. lenta* 2243 (2243) or *E. lenta* 15644 (15644) for 2 weeks. (A) Representative flow plots of Th17 (CD4+ IL-17a+ cells within the live CD3+ population) from the small intestinal lamina propria and the fold change of this population relative to the GF 1% arginine is quantified (B). Data are from 2 independent experiments (n=8-11). (C-G) SPF mice on a 1% or 3% arginine (Arg) diet were gavaged every other day with BHI, *E. lenta* 2243, or *E. lenta* 15644 for two weeks and then 2% DSS was administered and (C) disease scores were tracked for 7 days. \*\**P* < 0.01 two-way ANOVA with Sidak’s multiple comparison correction (1% vs. 3% 2243). (n=5-10). (D) Relative colon lengths of SPF gavaged with BHI, *E. lenta* 2243 or 1356 on a 1% or 3% Arg diet given 2% DSS. (E) Relative lipocalin in the colon content from the same mice as in (D). Data are from 2 independent experiments and set relative to the 15644 group on the 1% Arg diet which is set to 1. (F-G) Representative flow and percentages of IL-17a+ CD4+ cells within the live CD3+ population from the small intestinal lamina propria. \**P* < 0.05; \*\**P* < 0.01; \*\*\**P* < 0.001; \*\*\*\**P* < 0.0001 or stated (one-way ANOVA with Tukey multiple comparison test unless otherwise stated). Mean±SEM is displayed. Each point represents an individual mouse. See figs. S3 for gating strategies and S9 for related data.

The immunomodulatory effects of members of the human gut microbiota have long been appreciated and the microbiome field is now making efforts to explore the variety of mechanisms whereby these microbes influence a host’s immune response. Previous studies have identified bacteria capable of increasing Th17 levels and have implicated bacterial adherence to the gut epithelial in this activation (*15, 18, 28*). While we do not rule out that *E. lenta* can promote Th17 accumulation through an epithelial adhesion mechanism our results demonstrate that colonization is not necessary as conditioned media gavage is sufficient to induce intestinal Th17 cells. However, it is entirely possible that a single bacterium could increase Th17 cells through multiple mechanisms, as seen with SFB (*15, 29*). With recent studies profiling the immunomodulatory properties of a range of microbes spanning multiple phyla (*18, 30*), we have gained insight into how these associations could be leveraged for therapy (*31*). However, it is important to note that we may be missing many microbe-immune interactions due to the strain-level differences in immune modulation. For example, rat-derived SFB does not increase Th17 cells in mice, but mouse-derived SFB does (*15*). This concept also applies to bifidobacteria where *B. infantis* monocolonization increases Th17 cells but *B. longum* monocolonization does not (*18*); *B. infantis* is a subspecies of *B. longum*. These observations emphasize that strain-level variations have significant effects on host immune responses, and the current study outlines a generalizable strategy to determine the genes responsible. An important future area of study will be to investigate whether colonization with multiple inducers of Th17 cells results in a combinatorial effect on Th17 levels as this could inform as to whether these bacteria are inducing Th17 via similar or disparate mechanisms.

Cgr2 has been previously studied for its role in reducing the heart medication digoxin into a less active by-product, dihydrodigoxin (*12*). Digoxin is also a potent Th17 inhibitor and binds directly to the master transcription factor of Th17 cells, Rorγt (*26, 27*) while dihydrodigoxin is a less potent inhibitor (*32*). Yet digoxin is not normally present in the gastrointestinal tract and so the question arises whether Cgr2 is acting on another endogenous substrate. The substrate specificity of Cgr2 was recently explored and suggested that Cgr2 acts exclusively on the cardenolide class of molecules (*21*). Our working hypothesis is that Cgr2 acts on an endogenous Th17 inhibitory compound present both in mice and our BHI media that resembles digoxin (*33*) (fig. S10). Consistent with this hypothesis, a bile acid metabolite can inhibit Th17 cells by directly binding to Rorγt (*34*) and the levels of CD4+ IL-17a+ T cells and the MFI of Rorγt targets IL-17a and IL-17f were increased in *E. lenta* strain 2243 colonized mice but levels of Rorγt MFI and CD4+ Rorγt+ T cells were not. These findings suggest that *E. lenta* strain 2243 colonization may act by dampening the transcriptional regulation of Rorγt.

Diet is a key environmental factor that shapes the microbiota and microbial-mediated immune modulation. However, it is often difficult to assess the direct impact of dietary alterations on immune diseases from microbiota-mediated effects as a dietary alteration could have a range of effects directly on the host and microbiota that act in a synergistic or opposing fashion. Dietary arginine has multiple reported effects on mammalian cells (*35*), including iNOS-dependent suppression of inflammation (*36*), macrophage-dependent shifts in regulatory versus inflammatory skewing (*35, 37*), and T cell proliferation and cytokine production (*38, 39*). There are also multiple potential mechanisms through which arginine may interact with the gut microbiota, including diet-dependent shifts in gut microbial community structure and function (*10, 11*) or the microbial metabolism of arginine into polyamides (*40*), which have immunomodulatory functions and can alter microbial metabolism (*12*). Our results further emphasize that the effects of arginine on host phenotype is both concentration- and microbiome-dependent and provide a tractable model to continue to dissect the complex mechanisms that link dietary amino acids, the gut microbiota, and autoimmune disease.

## Acknowledgments

We acknowledge the UCSF gnotobiotic core (Jessie Turnbaugh, Kimberly Ly, Jolie Ma, Greg Ostolaza) for help with designing and execution of gnotobiotic experiments; Jody Baron and Jillian Jespersen for the *Rorc-/-* mice; Emily Balskus, Vayu Maini Rekdal, and Nitzan Koppel for the Cgr2 WT and Y333N plasmids and *R. erythropolis*; and Chunyu Zhao for assistance with MIDAS and IGGtools troubleshooting. We acknowledge the PFCC for assistance generating flow cytometry data (NIH P30 DK063720) and the UCSF Biorepository and Tissue Biomarker Technology Core for assistance with histology. We would also like to thank Kathy Lam, Tiffany Scharschmidt, and Mark Ansel for their critical reading and feedback on the manuscript. The model figure in S10 was made using BioRender.com software.

## Author contributions

Conceptualization, M.A. and P.J.T.; Investigation, M.A., Q.Y.A., R.R.N., V.U. A.E.B.; Resources, P.J.T.; Writing – Original Draft, M.A.; Writing – Review and Editing, M.A., Q.Y.A, R.R.N, V.U., P.J.T.; Supervision, P.J.T K.S.P.

## Data and materials availability

All data is available in the main text or the supplementary materials.

## Supplementary Materials

Materials and Methods

Figs. S1 to S10

Tables S1 to S2

Captions for Data S1 and S2

### Other Supplementary Materials for this manuscript include the following

Data S1 to S2 [DSS disease scores, Reagents]

