## Supplemental materials: methods, supplemental figures S1-10, and supplemental tables S1-2 for "A diet-dependent enzyme from the human gut microbiome promotes Th17 cell accumulation and colitis"

**This PDF file includes:**

Materials and Methods

Figs. S1 to S10

Tables S1 to S2

Captions for Data S1 and S2

**Other Supplementary Materials for this manuscript include the following:**

Data S1 to S2 [DSS disease scores, Reagents]

### Materials and Methods

#### Experimental Design

The objectives of these studies were to investigate whether and how a prevalent member of the human gut microbiota which is associated with autoimmune diseases, *Eggerthella lenta,* modulates T helper 17 (Th17) responses and mouse models of chronic inflammation and autoimmunity in a diet dependent manner. To investigate these questions, we utilized a combination of gnotobiotics, comparative genomics, *in vitro* Th17 skewing assays, mouse models of colitis, metagenomics, and bacterial genetics which are outlined below.

#### Mice

All mouse experiments were approved by the University of California San Francisco Institutional Animal Care and Use Committee. Housing conditions are specified (either gnotobiotic or SPF as described below). The mice were housed at temperatures ranging from 67-74°F and humidity ranging from 30-70%. No mice were involved in previous procedures before experiments were performed. Mice were assigned to groups to achieve similar age distribution between groups.

#### Gnotobiotic mouse studies

C57BL/6J mice (females and males, ages 6-16 weeks) were obtained from the University of California, San Francisco (UCSF) Gnotobiotics core facility (gnotobiotics.ucsf.edu) and housed in gnotobiotic isolators for the duration of each experiment (Class Biologically Clean) or were housed in Iso positive cages (Tecniplast). Mice were colonized via oral gavage with turbid mono-cultures of *E. lenta* ($10^9$ CFU/ml, 200 µl gavage) and colonization was confirmed via anaerobic culturing and or qPCR for an *E. lenta* specific marker (elnmrk1) (*19*, *21*). Mice were colonized for 2 weeks. For the heat-killed *E. lenta* preparation turbid *E. lenta* 2243 culture was incubated at 65°C for 15 min to kill bacteria.

<u>SPF mouse studies</u>

C57BL/6J mice (females or males, ages 6-10 weeks) were ordered from Jackson Labs. Mice were orally gavaged with turbid mono-cultures of *E. lenta* strains ($10^9$ CFU) every other day for 2 weeks and colonization was confirmed with qPCR for an *E. lenta* specific marker (elnmrk1) (*19, 21*). *Rorc-/-* (Rorctm2Litt) were generously provided by the Baron lab (UCSF). *IL-10-/-* mice were ordered from Jackson Labs and bred in house.

<u>DSS disease model</u>

For dextran sodium sulfate treatment (DSS) (Alfa Aesar, Cat no. 9011-18-1), single sexed mice (either all male or all female for both experimental and control groups) were given 2% DSS (w/v) *ad libitum* in their drinking water for 6-7 days after mice were colonized with *E. lenta* for 2 weeks prior to disease (either once for monocolonized mice or every other day gavage for SPF animals). SPF mice were additionally gavaged every other day throughout DSS treatment. Mice were monitored for disease progression and weighed daily. Gross signs of toxicity, including hematochezia and weight loss greater than 15% were monitored in this study and mice showing these signs were immediately euthanized. Stools were scored as follows: 0 = normal stool consistency, 1 = soft stool, 2 = blood in stool, 3 = bloody rectum, 4 = prolapsed rectum, 5 =

moribund/death (scoring based on (*41*)). For the SPF experiments with the 1% and 3% arginine diets, turbid *E. lenta* cultures (2243 and 15644) were spun down at 2000 rpm and resuspended in same amount of equilibrated fresh BHI media as *E. lenta* 2243 conditioned media is sufficient to promote Th17 accumulation in SPF mice.

*IL-10-/-* colitis model

For *IL-10-/-* colitis, mice were aged 6 to 10 weeks at the beginning of the experiment with an even distribution between groups (BHI or *E. lenta* strain 2243) and both males and females were used (with similar disease incidence seen between sexes). *IL-10-/-* mice were orally gavaged with 200 μl of turbid monocultures of *E. lenta* strains ($10^9$ CFU) 3 times a week in 3 independent experiments where gavage was carried out for 4, 6, or 10 weeks. Mice were weighed weekly and assessed for rectal prolapse incidence. Mice were sacrificed when they reached the endpoint (rectal prolapse) outlined in our animal protocol. Lipocalin levels in the colon content were observed via ELISA after 6 or 10 weeks of gavage of mice that remained in the study.

Diets

Custom diets with 1% (TD.170862) or 3% (TD.170863) Arg were purchased from Envigo. Otherwise, a chow diet (Lab Diet 5058) was used for SPF mice and an autoclaved chow diet (Lab Diet 5021) was used for the gnotobiotic mice. Diets used for gnotobiotic experiments were either autoclaved or irradiated and vacuum sealed to ensure sterility.

Human subjects

Consecutive patients from the UCSF Parnassus Rheumatology Clinic were screened for the presence of rheumatoid arthritis (RA) based on American College of Rheumatology criteria (*42*). RA patients were excluded if they had received prior therapy for RA with a disease-modifying anti-rheumatic drug (DMARD) or biologic therapy. Healthy controls were enrolled in the same clinic and were unrelated volunteer donors. After informed consent was signed, patients were provided with toilet hats (Collection Hat, Ability Building Center, NC0441080, Stool Collection Device), sample containers (Collection Vial, Fisher, NC9779954, Sarstedt Brown-Cap Vial) and swabs (Spectrum, 220135, BBL CultureSwab(TM)), cold packs (Fisher Scientific NC0515011) and pre-paid thermal envelopes (Polar Tech 116 Item # 116, Cool Barrier Bubble Economy Next Day) for home collection. Fresh feces were either immediately frozen at home or immediately shipped on frozen ice packs via USPS Priority Express overnight shipping. Samples were placed at -80°C upon receipt. The exclusion criteria applied to all groups were as follows: recent (<3 months prior) use of any antibiotic therapy, current extreme diet (*e.g.,* parenteral nutrition or macrobiotic diet), known inflammatory bowel disease, known history of malignancy, current consumption of probiotics, any gastrointestinal tract surgery leaving permanent residua (*e.g.*, gastrectomy, bariatric surgery, colectomy), or significant liver, renal, or peptic ulcer disease. This study was approved by UCSF Institutional Review Board (IRB). DNA was extracted using the MagBead ZymoBIOMICS 96 MagBead DNA Kit (D4302), see (*43*) for detailed protocol. Bead-beating was done with a Biospec Mini-Beadbeater-96 for 5 minutes.

Bacterial culturing

*E. lenta* strain information can be found in table S1. *E. lenta* strains were cultured at 37°C in an anaerobic chamber (Coy Laboratory Products) (2-5% $H_2$, 20% $CO_2$, balance $N_2$). Culture media was composed of brain heart infusion (BHI) media supplemented with *L*-cysteine-HCl (0.05%, w/v), hemin (5 µg/ml), arginine (1%), vitamin K (1 µg/ml), and resazurin (0.0001%, w/v) (BHI CHAVR). *E. lenta* strains were previously isolated and sequenced (*19*). *R. erythropolis* strain L88 was cultured in aerobic conditions at 30°C with shaking at 200 rpm in BHI CHAVR. *B. adolecentis* strain BD1 was isolated as described (*44*) and was cultured at 37°C in an anaerobic chamber in culture media comprising brain heart infusion (BHI) media supplemented with L-cysteine-HCl (0.05%, w/v), resazurin (0.0001%, w/v), hemin (5 µg/ml) and Vitamin K (1 µg/ml).

### Heterologous expression

Cardiac glycoside reductase 2 (*cgr2*) was heterologously expressed in *R. erythropolis* strain L88 as previously described (*21*). In short, competent *R. erythropolis* were electroporated [2.5 kV pulse (time constant ~4.8)] with pTipQC plasmid carrying the wild-type (WT) (Cgr2 WT) or Y333N mutated *cgr2* gene (Y333N Cgr2) and transformed cells were selected on Luria-Bertani (LB) chloramphenicol (17 µg/ml) plates. Liquid cultures were grown in BHI CHAVR with 34 µg/ml chloramphenicol to ~0.6 optical density (OD) and then treated with or without Thiostrepton (0.1 µg/ml) to induce expression. Conditioned media was harvested 48 hours later, centrifuged at 2500 rpm for 10 min to pellet the cells and debris then passed through a 0.2 µm syringe filter and used for cell culture assays. Expression induction was verified to be similar levels in the WT and Y333N plasmids via qRT-PCR for Cgr2 relative to dinB (a control for total *R. erythropolis* load) (*45*). For mouse experiments 200 µl *R. erythropolis* conditioned media or media control (BHI CHAVR with chloramphenicol (34 µg/ml) and thiostrepton (0.1 µg/ml) was gavaged every other day for 2 weeks from the *R. erythropolis* strains (WT Cgr2 or Y333N Cgr2). Liquid cultures were prepared as above (bacterial culturing).

Th17 skewing assay

Red blood cell (RBC) lysed mouse splenocytes from male or female C57BL/6J mice were filtered through a 40 µm filter and used for T cell isolation. T cells were isolated via Dynabeads untouched mouse CD4 isolation kit (ThermoFisher) according to kit specifications. Purity of the cells was assessed via flow cytometry for CD4+ cells and ranged from 90-95% CD4+ within the lymphocyte gate (fig. S8A). In a 96 well plate pre-coated with anti-CD3 (5 mg/ml, overnight 37°C), equal cell numbers were plated and were treated with bacterial conditioned media or unconditioned media controls with pH adjusted to 7 at a concentration of 5% or 7.5% volume/volume. At the same time Th17 skewing conditions were supplied (anti-CD28 (10 µg/ml), TGFβ (0.3 ng/ml), IL-6 (20 ng/ml), anti-IFNγ (2 mg/ml), anti-IL-4 (2 mg/ml) (*26*). Bacterial conditioned media was harvested from 48 hour stationary cultures where bacterial cells were pelleted (2500 rpm 10 min) and the supernatant was filtered through a 0.2 µm filter to exclude cells from the conditioned media preparation. Isolated CD4+ T cells were developed in Th17 skewing conditions with bacterial conditioned media present for 4 days at 37°C and then re-stimulated with PMA (50 ng/ml) and ionomycin (1000 ng/ml) overnight, then supernatants were harvested for IL-17a quantification via ELISA. For Supplemental Fig. 7, the same set up as above was performed but the conditioned media was added either at the time of skewing (pre) or after 4 days of skewing during overnight stimulation with PMA (Phorbol 12-myristate 13-acetate) and ionomycin (post). MTT (3-(4,5-dimethylthiazol-2-yl)-2,5-diphenyltetrazolium bromide) proliferation assay celltiter 96 non-radioactive cell proliferation assay (Promega G4000) was used to assay proliferation as per manufacturer's directions. Briefly, 15 µl of dye solution was added, the plate was incubated for 4 hours at 37°C, 100 µl stop solution was added and absorbance was immediately assessed with absorbance at 570 nm.

ELISAs

To measure the levels of secreted IL-17a from the Th17 cell culture assay we utilized the mouse IL-17a (homodimer) ELISA (ThermoFisher) according to the kit instructions, where 100 µl of cell culture media was loaded into ELISA. Raw values for IL-17a ELISAs are listed in table S2. To measure lipocalin levels in the colon contents 10 mg of colon content was resuspended in 200 µl of 0.1% tween20 in PBS, vortexed for 20 min at speed 8-9, spun for 10 min at 12,000 rpm to pellet debris. The supernatant was collected and used for a lipocalin ELISA (Mouse Lipocalin-2/NGAL DuoSet ELISA, R&D systems) according to the manufacturer's instructions, where 100 µl of the supernatant was loaded onto a coated ELISA plate. Absorbance for ELISA was measured at 450 nm and blank background signal was subtracted.

Lamina Propria Lymphocyte Isolation

Lamina propria lymphocytes (LPLs) were isolated with modifications of previously described methods (*46–48*). Briefly, small intestinal (SI) Peyer's patches were excised and colons and the lower ⅔ of the SI tissue were splayed longitudinally with mucus removed and stored in complete RPMI (10% fetal bovine serum, 100 units per ml penicillin and streptomycin, β-mercaptoethanol, glutamax, sodium pyruvate, hydroxyethyl piperazineethanesulfonic acid (HEPES) and non-essential amino acids). Media was removed by filtering through a 100 µM filter, and remaining tissue incubated in 1X Hank's Balanced Salt Solution (HBSS -without $Ca^{2+}$ and $Mg^{2+}$) containing 5 mM ethylenediaminetetraacetic acid (EDTA) and 1 mM DL-Dithiothreitol (DTT) for 45 min at 37°C on a shaker (200 rpm). Supernatant were filtered through a 100 µM filter, and remaining tissue was incubated for 45 min (colon) or 35 min (SI) at 37°C on a shaker in a solution containing 1X HBSS containing 5% (v/v) fetal bovine serum (GIBCO heat inactivated), 1 U/ml Dispase

(Sigma), 0.5 mg/ml Collagenase VIII (Sigma), and 20 μg/ml DNaseI (Sigma). The vortexed supernatant was filtered over a 40 μm cell strainer into 1X PBS. Cells were subjected to a Percoll (VWR) gradient (40%/80% [v/v] gradient) and spun at 2000 RPM for 20 min with no brake and no acceleration. Cells at the interface were collected, washed in 1X PBS and prepared for flow cytometry analysis as described in the next section.

Flow Cytometry

Lymphocytes were isolated from the colonic and SI lamina propria as described above. Spleen cells were prepped through gentle mashing with a syringe plunger. Spleen cells were treated with 1X RBC Lysis Buffer (Biolegend) to lyse and remove red blood cells. Surface staining for lymphocytes was done in staining buffer (1X HBSS (Corning) supplemented with 10 mM HEPES (Fisher Scientific), 2 mM EDTA (Invitrogen), and 0.5% (v/v) fetal bovine serum (heat inactivated - 30 minutes at 56°C with mixing)) for 20 min at 4°C. Cells were then washed twice in supplemented 1X HBSS and enumerated via flow cytometry. The following antibodies were used: anti-CD3 (17A2, Fisher Scientific), anti-TCRβ (H57-597, Biolegend), anti-CD4 (GK1.5, Biolegend), and live/dead staining was performed using LIVE/DEAD Fixable Dead Cell Stain Kit (Life Technologies). For intracellular staining, cells were first stimulated with ionomycin (1000 ng/ml), PMA (50 ng/ml), and Golgi Plug (1 μl/sample) (BD Bioscience) 4-6 hours or overnight at 37°C. Alternatively, cells were stimulated with a cell stimulation cocktail (Fisher Scientific) containing PMA and ionomycin according to the manufacturer's instructions, and Golgi plug was added. Cells were surface stained, washed, and then fixed/permeabilized in 100 μl fixation and permeabilization (Perm) buffer (BD Bioscience). Cells were washed twice in Perm/Wash buffer (BD Bioscience) and then stained for intracellular cytokines with the following antibodies: anti-

IFNγ (XMG1.2, Fisher Scientific), anti-IL17a (ebio17B7, Invitrogen), Rorγt (B2D, ebioscience). Cells were washed twice in Perm/Wash buffer and then placed in staining buffer for flow cytometry analysis. Gating cell populations was done using isotype and single stain controls. Gating strategies and which figures they correspond to are outlined in fig. S2. Live/dead staining or forward or side scatter (FCS SSC) occasionally revealed a high death incidence in some lamina propria lymphocyte isolations after PMA and ionomycin stimulation. These samples were removed from our analysis of immune cells as they had high levels of non-specific background fluorescence and too few cells to analyze. The flow cytometry data were collected with a BD LSR Fortessa and analyzed with FlowJo software (version 10.6.1) and a list of all antibodies are included in Data S2.

Histology

1 cm sections of the distal colon were collected for histological processing from DSS treated mice. Samples were fixed in formalin for 24 hours and then stored in 70% ethanol. Samples were processed by the UCSF Biorepository and Tissue Biomarker Technology Core. Tissues were embedded in wax and 4 µm cross-sections were hematoxylin and eosin (H&E) stained.

Comparative genomics

To identify shared genetic regions of IL-17a inducing *E. lenta* strains which were also excluded from non-IL-17 inducing *E. lenta* strains we utilized ElenMatchR (github.com/turnbaughlab/ElenMatchR) (*19*). Briefly, gene presence/absence is used as the input variable for a random forest classifier against user-provided phenotypes or traits, in this case induction or non-induction of IL-17a by *E. lenta* strains. With these classifications, we performed comparative genomics using ElenMatchR (80% coverage and 80% minimum identity 3 replications of the random forest analysis and 1000 trees generated in the model) to determine genomic regions shared between the 4 inducing strains and absent from the 6 non-inducing strains.

RNA/DNA isolation and qRT-PCR

Ileal segments from GF or *E. lenta* monocolonized mice were harvested and RNA was extracted with Direct-zol RNA MiniPrep kit (Zymo) according to manufacturer instructions for tissue RNA isolation. Tissue was homogenized with a mortar and pestle and DNase treatment was performed on columns. RT-PCR was performed with iScript (BioRad) using ~300 ng RNA and qPCR using SYBR select. DNA was extracted from mouse and human fecal and cecal samples to quantify levels of *E. lenta* or Cgr2 with the ZymoBIOMICS 96 MagBead DNA Kit according to manufacturer's instructions. Briefly, samples were weighed out (~50 mg), lysed, disrupted for 5 minutes in Biospec beadbeater, washed, and eluted in 50 µl (*43*). For quantification of Cgr2 from human fecal samples qPCR was performed with ~10 ng gDNA using SYBR select and levels were normalized to input mg of fecal content. For *E. lenta* quantification DNA was extracted from mouse cecal content (~50 mg) and *E. lenta* specific primers (elnmrk1) were used with SYBR select with 3 µl of gDNA. Primers are listed in Data S2.

Metagenomic data analysis

Metagenomic analysis was performed on samples from an inflammatory bowel disease (IBD) metagenomic survey of Chinese and Spanish (MetaHIT) patients (*22–24*). Shotgun metagenomic reads were analyzed with an implementation of Metagenomic Intra-Species Diversity Analysis Subcommands (MIDAS) (*49*) designed for a Unified Human Gastrointestinal Genome (UHGG) collection of 286,997 isolate genomes and metagenome assembled genomes from the human gut environment. Presence of *E. lenta* in a given sample was established by 15 or more reads mapping to *E. lenta* single-copy universal genes (HS-BLASTN) (*50*), with greater than 70% coverage. Because species read counts are by nature compositional, species relative abundance between sample types was determined by centered-log ratio (clr) transformation of species read counts, and subsequent Wilcoxon rank test for significance using ALDEx2 (*51*), run with 128 Monte Carlo samples of the Dirichlet distribution for each sample (*52–54*). In addition, rank relative abundance of *E. lenta* was assessed by ranking all species by their median relative abundance in a given disease type.

Statistical Analysis

Statistical analyses were performed using GraphPad Prism software (Version 8). ANOVA with Tukey's multiple comparison test was used for the parametric analysis of variance between groups, and unpaired Welch's *t* tests were used for pairwise comparisons. The Mann-Whitney nonparametric test was used to compare *cgr2* levels in RA patients and healthy stool. Two-way ANOVAs with Sidak's multiple comparisons test were used to compare disease scores and weights over time. Outliers were removed as determined by ROUT (Q = 10%) or Grubb's (alpha = 0.2) methods. Numbers (*n*) are stated and displayed as individual points on plots.

**Supplementary Figures:**

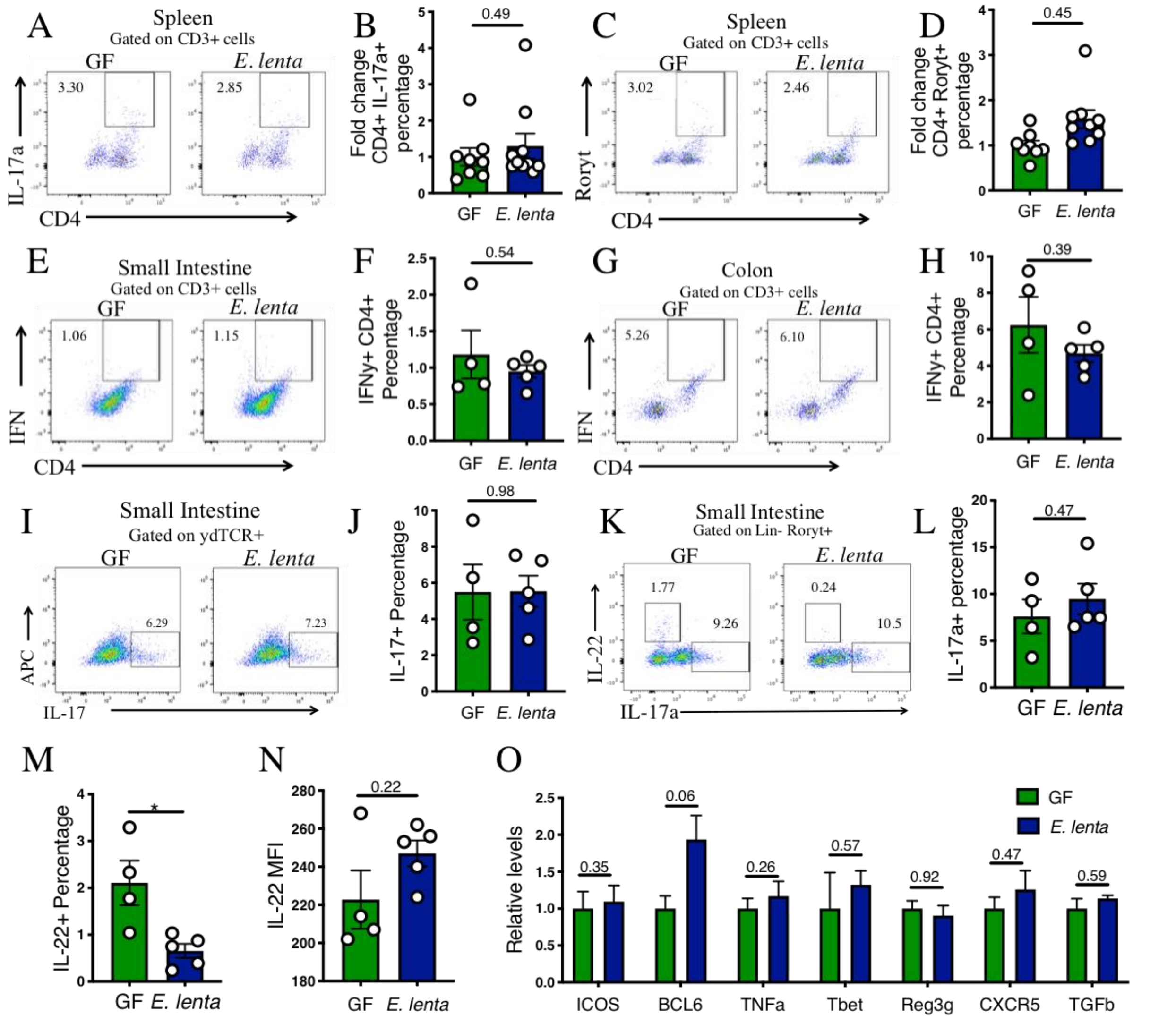


**Fig. S1. Splenic Th17 cell frequencies, Th1, IL-17+ γδ T cells and group 3 innate lymphoid cells (ILC3), and immune gene expression in germ-free (GF) and *E. lenta* monocolonized mice.** (A-D) GF or *E. lenta* strain 2243 monocolonized mice (A) splenic CD4+ IL-17a+ gated on CD3+ cells and (B) fold change. (C) Splenic CD4+ Rorγt+ gating on CD3+ cells representative flow and (D) fold change. Data are from 2 independent experiments (n=8-10). (E-H) Representative Th1 cells and fold change (IFNγ+ CD4+ gated on CD3+ cells) in the (E-F) small intestinal and (G-H) colonic lamina propria from GF mice or mice monocolonized with *E. lenta* strain 2243. Data are from 1 independent experiment (n=4-5). (I-J) IL-17a+ cells within the γδTCR+ population representative flow and quantification to the right from GF mice or mice monocolinzed with *E. lenta* strain 2243. (K-M) Lineage- Rorγt+ IL17a+ and Lineage- Rorγt+ IL-22+ representative flow and frequencies (L) of the IL-17a+ and (M) IL-22+ cells from GF mice or mice monocolinzed with *E. lenta* strain 2243. Data are from 1 experiment (n=4-5). (N) IL-22 MFI gated on total lymphocytes (n=4-5). (O) RT-qPCR panel of immune genes from GF or *E. lenta* monocolonized ileal samples (n=4). Data are relative to GF. $*P < 0.05$; or stated Welch's t-test. Mean±SEM is displayed. Each dot represents an individual mouse.

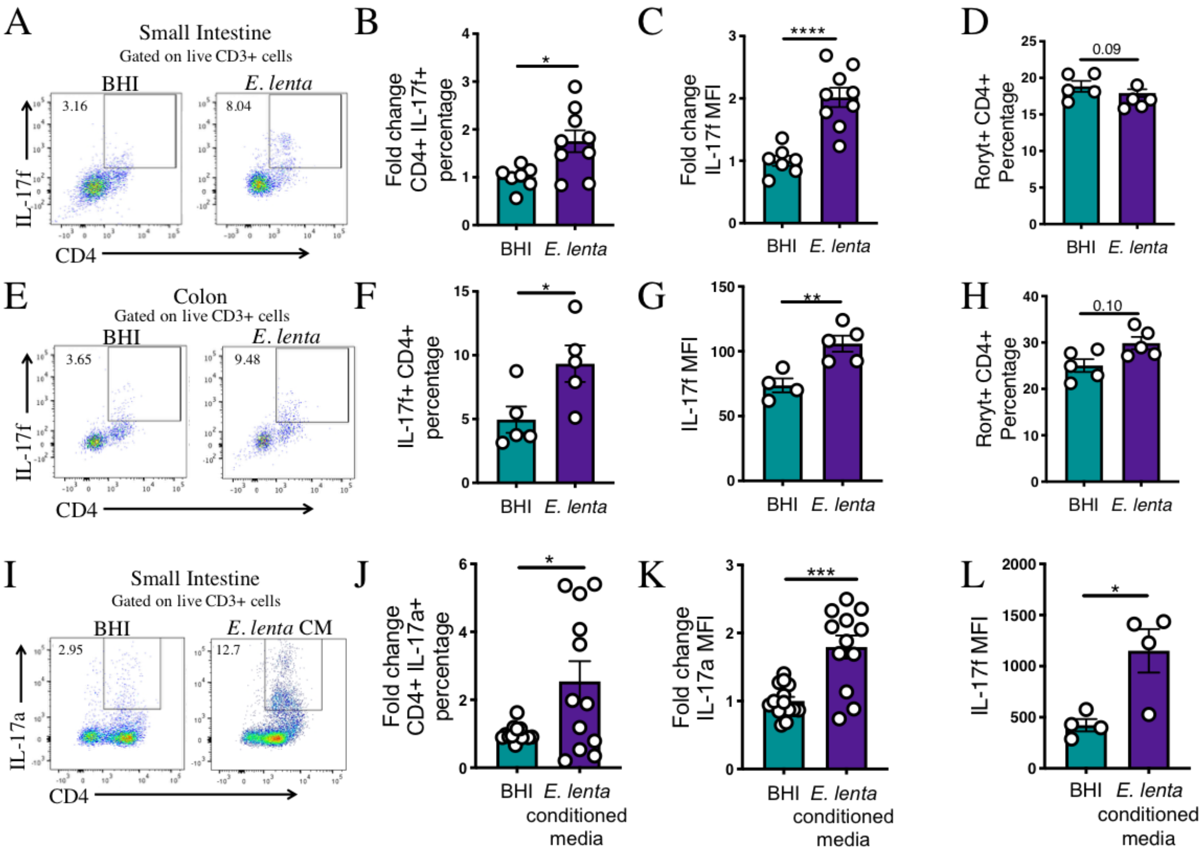


**Fig. S2. Intestinal CD4+ IL-17f and IL-17a levels in SPF mice gavaged with BHI, *E. lenta*, or *E. lenta* conditioned media.** (A-H) C57BL/6J SPF mice were gavaged every other day for 2 weeks with BHI media control or *E. lenta* strain 2243. (A-C) Representative flow plots of small intestinal lamina propria IL-17f+ CD4+ cells within the live CD3+ population and fold change of this population and IL-17f MFI are plotted to the right (n=7-9). Data are from two independent experiments. (D) Percentages of small intestinal CD4+ Rorγt+ within the live CD3+ population (n=5). Data are from one experiment. (E-F) Representative flow plots of colonic lamina propria IL-17f+ CD4+ cells within the CD3+ population and fold change of this population and IL-17f MFI are plotted to the right (n=5). (H) Percentages of colonic CD4+ Rorγt+ within the CD3+ population (n=5). Data are from one experiment. (I-L) C57BL/6J SPF mice were gavaged with BHI or *E. lenta* conditioned media (CM) every other day for 2 weeks. (I-K) Levels of CD4+ IL-

17a+ cells within the live CD3+ population and IL-17a MFI were quantified via flow cytometry in the small intestinal lamina propria. Data are from three independent experiments (n=14). (L) IL-17f MFI (n=4) from one independent experiment. Each point represents an individual mouse and the mean±SEM is plotted. *$P$ < 0.05; **$P$ < 0.01; ***$P$ < 0.001; ****$P$ < 0.0001 or listed, Welch's t-test.

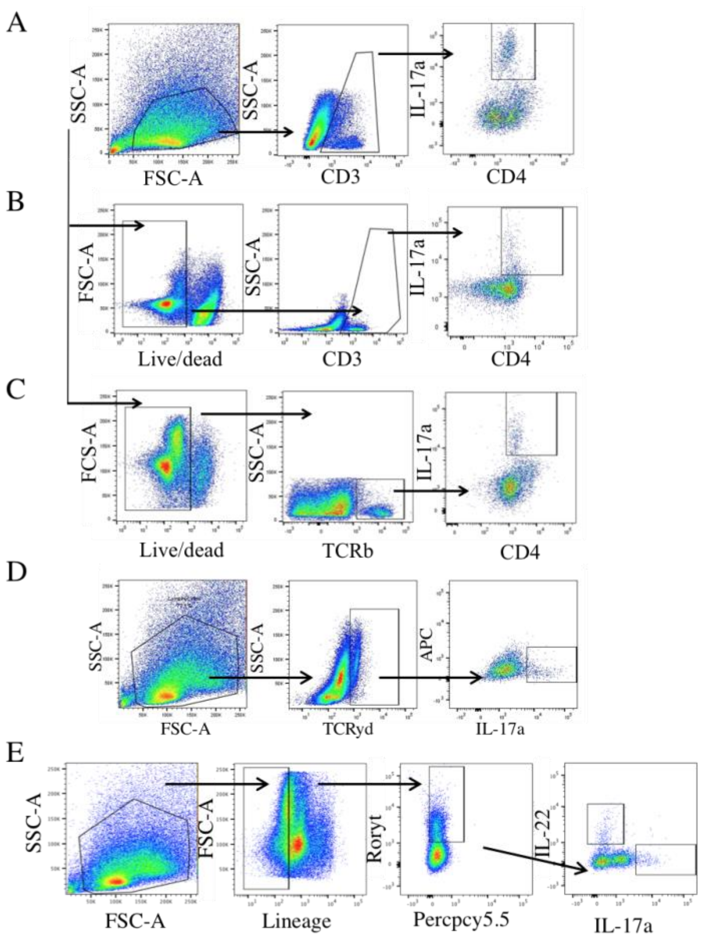


**Fig. S3. Gating strategies.** (A) Gating strategy pertaining to figures 1A, 1D, 1G, 1J, 3H and supplemental figures 1A, 1C, and 5B. (B) Gating strategy pertaining to figures 4D, 5A, 5F and supplemental figures 2A, 2E, 2I, 5E, 5G, 6A and 6E. (C) Gating strategy pertaining to figures 4D and C. (D) Gating strategy pertaining to Supplemental Figure 1I. (E) Gating strategy pertaining to Supplemental Figure 1K.

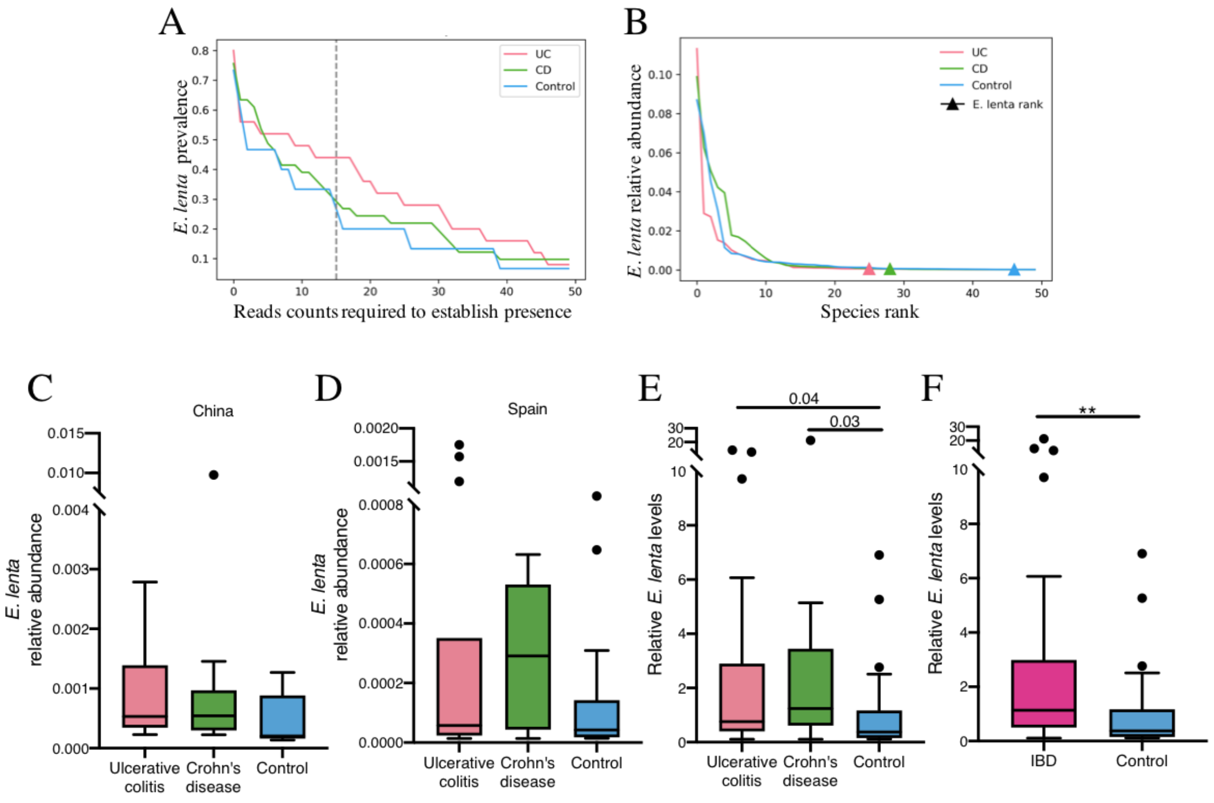


**Fig. S4. *E. lenta* is enriched in inflammatory bowel disease patients relative to healthy controls.** (A-C) Prevalence and abundance of *E. lenta* was assessed using Metagenomic Intra-Species Diversity Analysis Subcommands (MIDAS) to map shotgun metagenomic reads to a Unified Human Gastrointestinal Genome (UHGG) collection of isolate genomes and metagenome assembled genomes from the human gut environment (*55–57*). The analyzed shotgun reads were obtained from an inflammatory bowel disease (IBD) metagenomic study performed in China, that included individuals with ulcerative colitis (UC) (n = 25), Crohn's disease (CD) (n = 41), and controls (n = 15) (*22*). Percent of human IBD samples that are *E. lenta* positive versus mapped read counts required to establish presence *E. lenta* in a sample. A read count cutoff (gray dashed line) of 15 was chosen because *E. lenta* presence is based on reads mapping to any of 15 single copy universal genes. (B) Rank relative abundance profiles for each sample type were determined by ranking species based on median relative abundance. *E. lenta* rank is indicated by triangle, for each disease label. (C) *E. lenta* relative abundance by sample type in China cohort within *E. lenta* positive samples. Normalized species reads within a sample by the total number of reads across all species in the sample. (D) *E. lenta* relative abundance within *E. lenta* positive samples by sample type in Spain cohort ulcerative colitis (n = 27), Crohn's disease (n = 12), and controls (n = 85) as is C. (E) Combined levels of *E. lenta* set relative to control from the Spain and China cohorts. Listed *P* values are Kruskal-Wallis with Benjamini-Hochberg correction. (F) Combined levels of *E. lenta* set relative to control from the Spain and China cohorts with both Crohn's disease and ulcerative colitis patients combined into the IBD category. **$P < 0.01$ Mann-Whitney test.

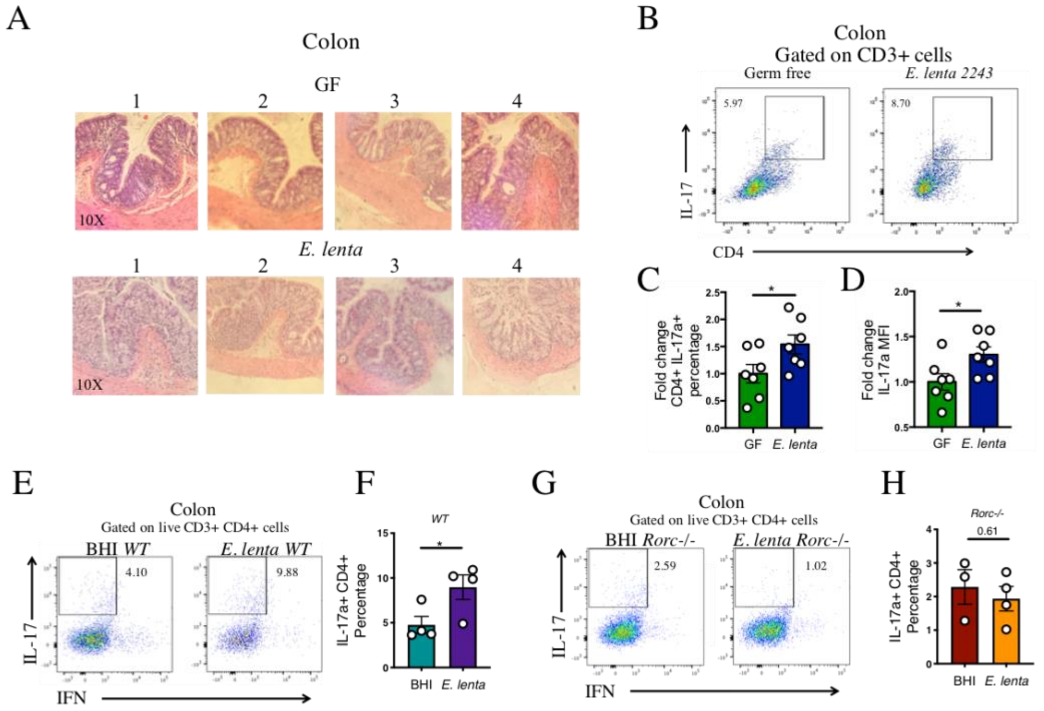


**Fig. S5. Colon histology and Th17 accumulation in *E. lenta* colonized DSS treated mice.** (A) Histology from GF or *E. lenta* monocolonized mice on 2% DSS for 6 days (n=4) (B-D) Representative flow of colonic lamina propria CD4+ IL-17a+ within the CD3+ gate from germ free (GF) or *E. lenta* 2234 monocolonized mice treated with 2% DSS for 6 days, (C) frequencies and (D) IL-17a MFI are quantified below from a combination of 2 independent experiments (n=7). **P* < 0.05, ***P* < 0.01 Welch's t-test (E-H) *WT* and *Rorc-/-* SPF mice were gavaged with BHI or *E. lenta* strain 2243 every other day for 2 weeks and then treated with 2% DSS while gavages continued. On day 7 of DSS treatment colonic lamina propria IL-17a+ cells within the live CD3+ CD4+ population were quantified via flow cytometry and representative flow plots are displayed and frequencies are displayed (n=3-4). **P* < 0.05 or stated (Welch's t-test). Mean±SEM is displayed. Each dot represents an individual mouse.

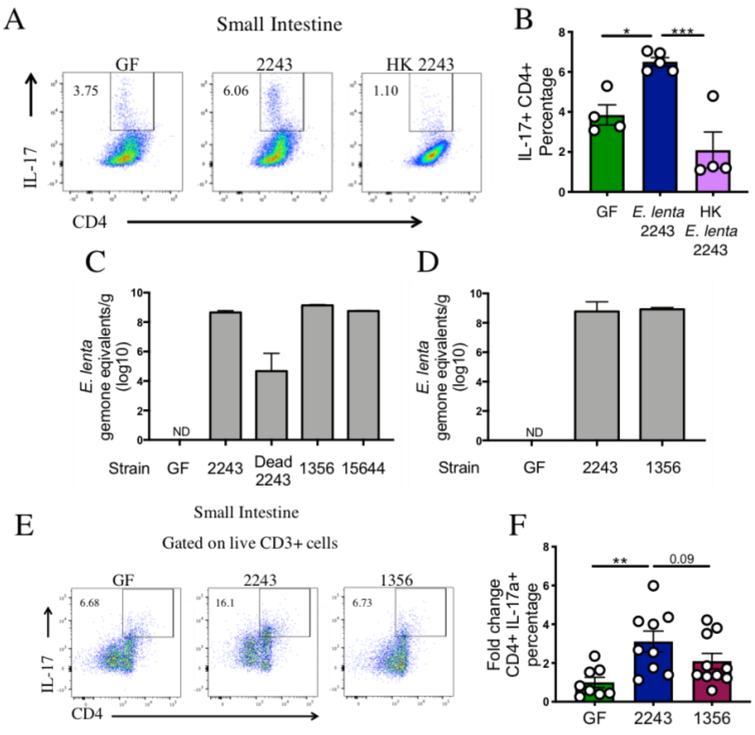


**Fig. S6. Th17 cell frequency in *E. lenta* strain gnotobiotic experiments and colonization levels.** (A) Representative flow of CD4+ IL-17a+ population with (B) frequencies quantified from GF mice or mice monocolonized with *E. lenta* 2243, or heat killed *E. lenta* 2243 (HK 2243) in the small intestinal lamina propria. (n=4-5) (C and D) Quantification of *E. lenta* levels in gnotobiotic experiments from *E. lenta* strain monoclonization experiments and (D) *E. lenta* stain monocolonization experiment with DSS treatment. (E-F) Th17 levels were measured in the small intestinal lamina propria and representative flow plots of IL-17a+ cells within the CD3+ population and fold change relative to the GF control are quantified to the right. Data are from two independent experiments (n=8-10). *$P < 0.05$; **$P < 0.01$; ***$P < 0.001$; or listed (one-way ANOVA with Tukey multiple comparison test). Mean±SEM is displayed. Each dot represents an individual mouse.

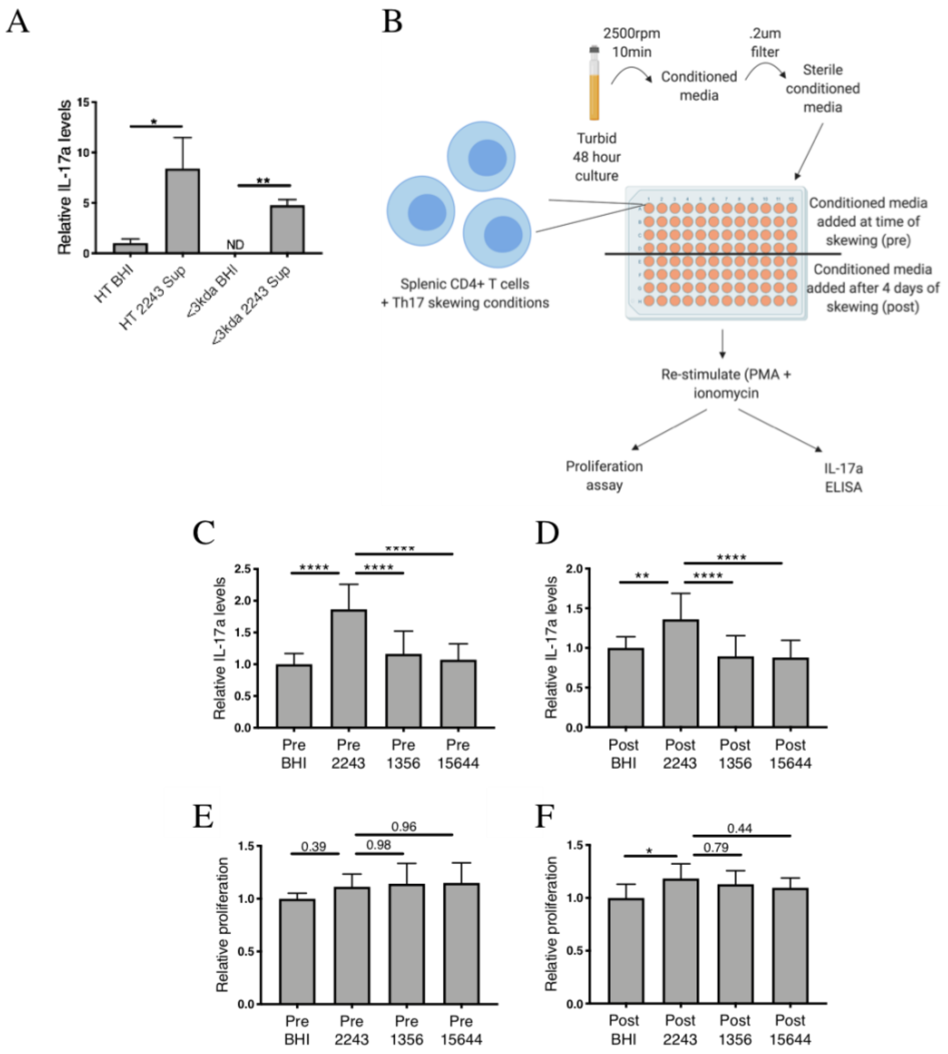


**Fig. S7. *E. lenta* strain 2243 conditioned media alters IL-17a production**. (A) Isolated splenic CD4+ T cells were treated with Th17 skewing conditions and conditioned media from *E. lenta 2243* heat treated at 95°C for 10 min. (HT 2243 Sup) or passed through a 3kda filter (<3kda 2243 Sup) with the same performed on BHI (HT BHI or <3kda BHI) media control. Relative levels of IL-17a are shown as measured by ELISA and are relative to the HT BHI group. $*P < 0.05$; $**P < 0.01$, Welch's t tests (n=4-8). Data are from 2 independent experiments. (B) Experimental design for testing whether conditioned media from *E. lenta* strain 2243, 1356, and 15644 impacts differentiation, proliferation, or expression of IL-17a. (C) IL-17a levels as measured by ELISA when conditioned media or controls are added when Th17 skewing conditions were applied (Pre). (D) IL-17a levels when conditioned media was added after cells were already skewed to Th17 fate (Post). Data are from two independent experiments (n=14). (E-F) Relative proliferation levels as quantified by the MTT assay from the "pre" and "post" scenarios described above. Data are from two independent experiments (n=9). $^{*}P < 0.05$; $^{**}P < 0.01$; $^{****}P < 0.0001$; or listed (one-way ANOVA with Tukey multiple comparison test). Mean±SEM is displayed.

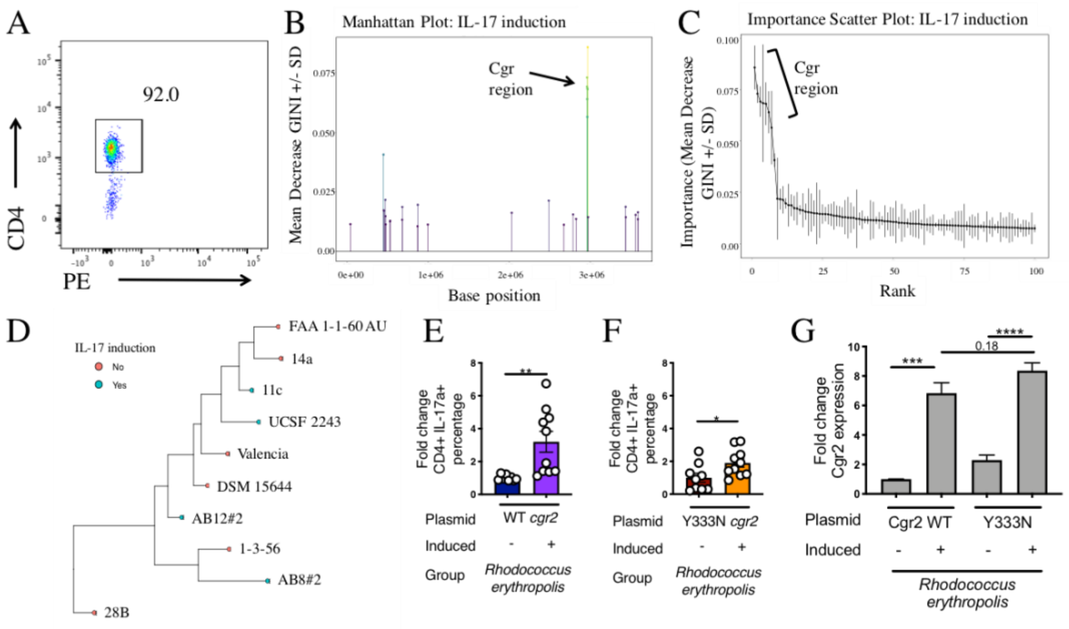


**Fig. S8. ElenMatchR analysis from *E. lenta* strain's Th17 induction *in vitro* experiments.** (A) Representative flow plot of CD4+ cells isolated from the spleen via Dynabeads untouched mouse CD4 isolation kit. (B) Manhattan plot of genome associations with IL-17a induction from our in vitro screening of *E. lenta* stains (Fig. 4B). (C) Importance scatter plot of genomic associations with IL-17a induction. For B and C the *cgr* region is highlighted. (D) Phylogenetic tree of *E. lenta* strains screened in the Th17 *in vitro* screen with IL-17a induction noted yes or no. Plots were produced by ElenmatchR (*19*, *58*). (E-F) C57BL/6J SPF mice were gavaged with conditioned media from *R. erythropolis* strains with induced (+Thiostrepton) or uninduced (-Thiostrepton) expression of WT Cgr2, Y333N Cgr2, or a BHI media control (with Thiostrepton). Fold change of CD4+ IL-17a+ cells within the live CD3+ populations (n=10). Levels are relative to the matching uninduced control (-thiostrepton). *$P < 0.05$; **$P < 0.01$, Welch's t tests (G) Relative levels of Cgr2 induction in *R. erythropolis* with heterologous expression of WT or Y333N mutated Cgr2 (relative to WT Cgr2 which is set to 1). ***$P < 0.001$; ****$P < 0.0001$; or listed (one-way ANOVA with Tukey multiple comparison test). Mean±SEM is displayed.

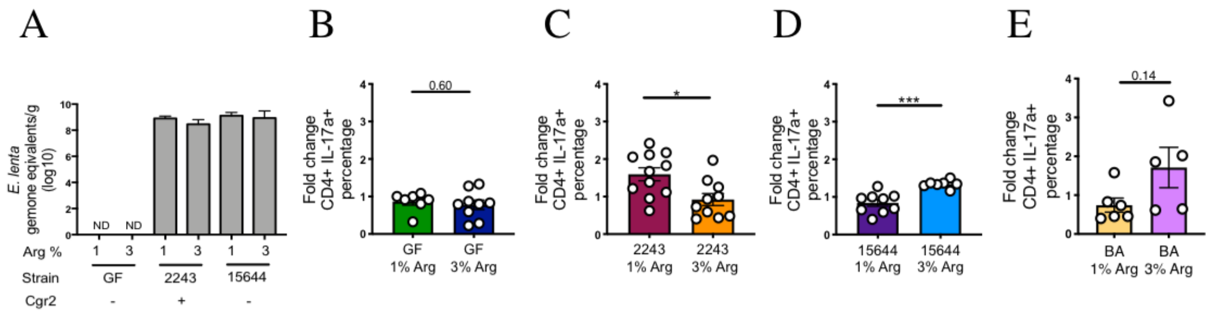


**Fig. S9. Arginine does not impact *E. lenta* colonization levels and has different effects on Th17 cell levels depending on colonization state.** (A) *E. lenta* colonization levels in GF, *E. lenta* 2243 or 1356 monocolonized mice on a 1% or 3% Arg diet as determined via qPCR. (B-E) Fold change in Th17 levels in the small intestine of (B) GF, (C) *E. lenta* 2243, (D) *E. lenta* 15644, or (E) *B. adolescentis* (BA) on a 1% or 3% arginine (Arg) diet. Levels are relative to the GF group on a 1% arginine diet which is set to 1. All scales are the same for comparison of magnitude. *P* values are stated or, *$P < 0.05$, ***$P < 0.001$ Welch's t-tests. Mean±SEM is displayed. Data are from two independent experiments except for the *B. adolescentis* groups which are from one independent experiment.

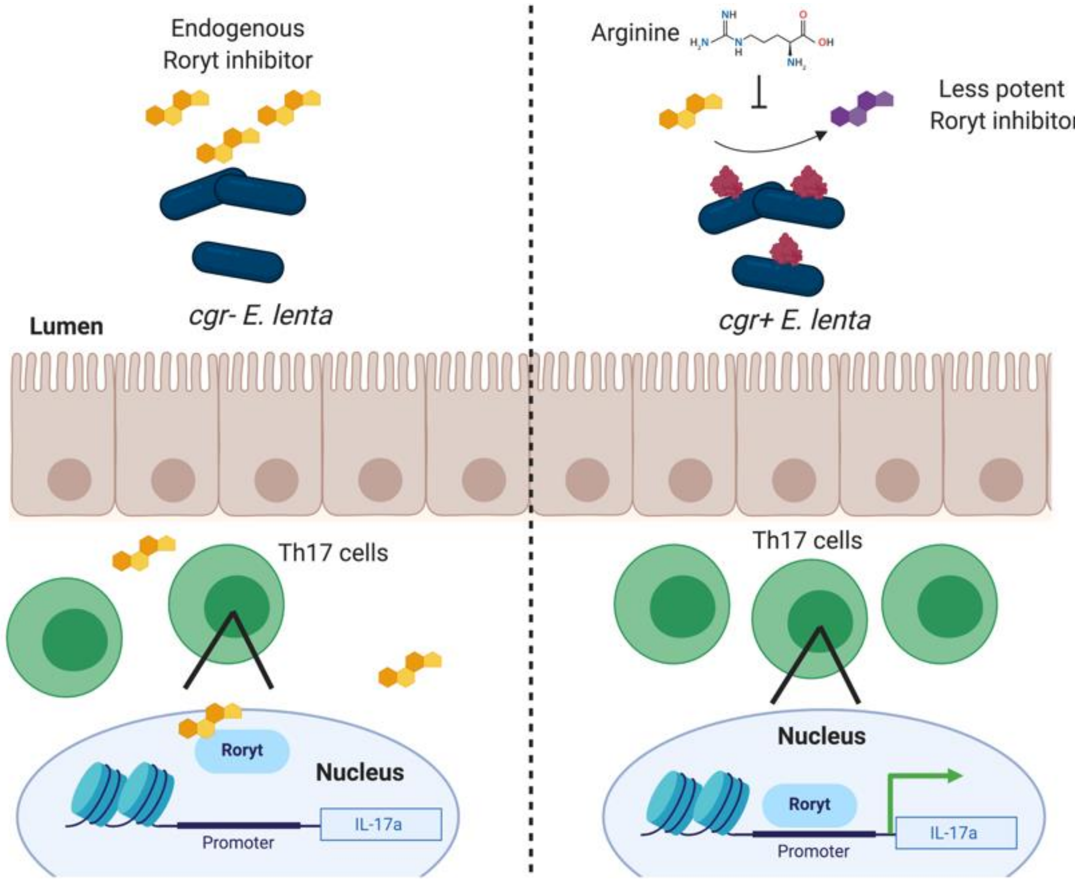


**Fig. S10. Working model.** Our proposed model of a mechanism for *E. lenta* mediated IL-17a elevation where high arginine levels inhibit the metabolism of an endogenous Rorγt inhibitor into a less potent form by *cgr+ E. lenta* resulting in increased transcription of Rorγt targets such as IL-17a.

**Supplementary Tables:**

**Table S1. Strain metadata.** For more information please refer to the original characterization of this strain collection (*19*).

| **Strain** | **Abbv. ID** | **Bioproject** | **BioSample** | **Genome accession** |
|---|---|---|---|---|
| *Eggerthella lenta* 14A | 14a | PRJNA412637 | SAMN08365961 | PPUR00000000 |
| *Eggerthella lenta* 28B | 28b | PRJNA412637 | SAMN08365965 | PPUN00000000 |
| *Eggerthella lenta* DSM 15644 | 15644 | PRJNA412637 | SAMN08365977 | PPUB00000000 |
| *Eggerthella lenta* FAA 1-1-60AU | 1-1-60AU | PRJNA412637 | SAMN08365980 | PPTY00000000 |
| *Eggerthella lenta* FAA 1-3-56 | 1356 | PRJNA412637 | SAMN02463797 | GCA_000185625.1 |
| *Eggerthella lenta Valencia* | Valencia | PRJNA412637 | SAMN08365983 | PPTV00000000 |
| *Eggerthella lenta* 11C | 11c | PRJNA412637 | SAMN08365960 | PPUS00000000 |
| *Eggerthella lenta* AB12 #2 | AB12 | PRJNA412637 | SAMN08365968 | PPUK00000000 |
| *Eggerthella lenta* AB8 #2 | AB8 #2 | PRJNA412637 | SAMN08365969 | PPUJ00000000 |
| *Eggerthella lenta* UCSF 2243 | 2243 | PRJNA412637 | SAMN08365979 | PPTZ00000000 |

**Table S2. IL-17a ELISA levels.** Levels of IL-17a in pg/ml as measured via ELISA from *in vitro* Th17 skewing assays. NS denotes no treatment.

| Experiment 1 | | | | | Experiment 2 | | | | |
|---|---|---|---|---|---|---|---|---|---|
| **NS** | **BHI** | **2243** | **1356** | **15644** | **NS** | **BHI** | **2243** | **1356** | **15644** |
| 227.5 | 143.13 | 261.88 | 155 | 193.75 | 24.84 | 9.42 | 28.03 | 20.56 | 17.04 |
| 220 | 118.13 | 233.13 | 167.5 | 163.13 | 21.73 | 8.22 | 30 | 16.36 | 8.57 |
| 167.5 | 113.75 | 243.75 | 126.88 | 160.63 | 22.97 | 8.26 | 27.61 | 6.64 | 9.32 |
| 241.25 | 116.88 | 265.63 | 125 | 157.5 | 25.1 | 2.17 | 30.84 | | 8.52 |

| Experiment 3 | | | | | | | |
|---|---|---|---|---|---|---|---|
| **NS** | **BHI** | **2243** | **1356** | **15644** | **11c** | **Valencia** | **14A** |
| 49.38 | 60.63 | 130.63 | 3.13 | 13.13 | 73.13 | 65.63 | 4.38 |
| 124.38 | 143.75 | 171.25 | 4.38 | 20 | 129.38 | 48.13 | 6.88 |
| 108.13 | 148.13 | 145.63 | 4.38 | 18.75 | 128.75 | 36.25 | 4.38 |
| 45.63 | 71.25 | 120.63 | 5.63 | 33.13 | 173.75 | 67.5 | 7.5 |

| Experiment 4 | | | | | | | | | |
|---|---|---|---|---|---|---|---|---|---|
| **NS** | **BHI** | **2243** | **1356** | **15644** | **11c** | **22c** | **14a** | **AB8#2** | **AB12#2** |
| 49.36 | 13.71 | 92.84 | 44.58 | 37.62 | 119.8 | 66.32 | 48.06 | 112.41 | 159.36 |
| 35.45 | 11.1 | 84.14 | 38.49 | 21.54 | 68.49 | 54.14 | 37.62 | 103.27 | 121.54 |
| 43.27 | 16.75 | 88.49 | 46.32 | 27.62 | 79.36 | 58.93 | 34.58 | 89.36 | 118.06 |
| 27.62 | 12.41 | 82.41 | 45.01 | 26.75 | 68.06 | 41.54 | 25.45 | 80.23 | 117.19 |
| 46.75 | 11.54 | 100.23 | 40.67 | 21.1 | 78.06 | 49.8 | 32.41 | 89.8 | 118.06 |
| 23.27 | 35.45 | 95.88 | 44.58 | 21.97 | 111.1 | 51.54 | 25.45 | 102.84 | 117.19 |
| 25.88 | 14.58 | 83.71 | 45.88 | 30.23 | 92.84 | 79.36 | 33.71 | 99.36 | 108.06 |
| 28.93 | 11.1 | 63.71 | 43.71 | 24.14 | 93.71 | 35.45 | 35.88 | 115.01 | 121.97 |

**Supplemental Data Captions:**

**Data S1. DSS disease scores.** Raw disease scores of all DSS experiments.

**Data S2. Reagents.** A list of reagents used in this study.
